# Structure and evolution of Photosystem I in the early-branching cyanobacterium *Anthocerotibacter panamensis*

**DOI:** 10.1101/2024.10.31.621444

**Authors:** Han-Wei Jiang, Christopher J. Gisriel, Tanai Cardona, David A. Flesher, Gary W. Brudvig, Ming-Yang Ho

**Affiliations:** Department of Life Science, National Taiwan University, Taipei, Taiwan; Department of Chemistry, Yale University, New Haven, CT, USA; Department of Biochemistry, University of Wisconsin-Madison, Madison, WI, USA; School of Biological and Behavioural Sciences, Queen Mary University of London, London, United Kingdom; Department of Molecular Biophysics and Biochemistry, Yale University, New Haven, CT, USA; Institute of Plant Biology, National Taiwan University, Taipei, Taiwan

**Author notes:** These authors contributed equally to this work (Han-Wei Jiang, Christopher J. Gisriel, Tanai Cardona).

**Keywords:** Structural, biochemical, and evolutionary analyses of PSI in a thylakoid-free cyanobacterium, *Anthocerotibacter panamensis*

## Abstract

Thylakoid-free cyanobacteria are thought to preserve ancestral traits of early-evolving organisms capable of oxygenic photosynthesis. However, and until recently, photosynthesis studies in thylakoid-free cyanobacteria were only possible in the model strain Gloeobacter violaceus. Here, we report the isolation, biochemical characterization, cryo-EM structure, and phylogenetic analysis of photosystem I from a newly-discovered thylakoid-free cyanobacterium, Anthocerotibacter panamensis, a distant relative of the genus Gloeobacter. We find that A. panamensis photosystem I exhibits a distinct carotenoid composition and has one conserved low-energy chlorophyll site, which was lost in G. violaceus. These features explain the capacity of A. panamensis to grow under high light intensity, unlike other Gloeobacteria. Furthermore, we find that, while at the sequence level photosystem I in thylakoid-free cyanobacteria has changed to a degree comparable to that of other strains, its subunit composition and oligomeric form might be identical to that of the most recent common ancestor of cyanobacteria.

## Introduction

Oxygenic photosynthesis, occurring in cyanobacteria, algae, and plants, powers the biosphere by converting light into chemical energy and produces molecular oxygen, thus sustaining aerobic life on Earth (*1*). This process occurs in two multi-subunit pigment-protein complexes, photosystem I (PSI) and photosystem II (PSII) (*2, 3*). Central to oxygenic photosynthesis, PSI is responsible for light harvesting, charge separation, and electron transfer, leading to the reduction of NADP^+^ to NADPH, which is essential for CO2 fixation. The PSI complex comprises 10 to 12 protein subunits and various cofactors, including chlorophylls (Chls), carotenoids, quinones, and iron-sulfur clusters (*3, 4*). Most of these cofactors are well-conserved in the PSI cores of oxygenic phototrophs, although the oligomerization states and subunit compositions of PSI differ among species (*5*).

Low-energy Chls that absorb wavelengths longer than 700 nm are a distinguishing characteristic of PSI. This is interesting because the primary electron donor in PSI, a pair of Chl molecules called P700, requires energy equivalent to 700 nm to achieve charge separation (*6*). The low-energy Chls, which often form dimers and higher-order aggregates, are believed to deliver energy uphill to P700 (*7*). They are especially important when light with wavelengths less than 700 nm is limited in the environment (e.g., some shaded environments) and they participate in photoprotection by dissipating energy when P700 is oxidized (*8–11*). Identifying the positions of low-energy Chls bound to PSI is challenging due to the large number of Chls coordinated by the complex (*3, 4*). Despite extensive studies, the precise locations of these low-energy Chls in PSI have been challenging to elucidate.

A recent study using cryogenic electron microscopy (cryo-EM) presented a high-resolution structure of the trimeric PSI complex from an early-diverging thylakoid-free cyanobacterium in the Gloeobacteria family, *Gloeobacter violaceus* PCC 7421

(hereafter *G. violaceus*) (*12*). Structural comparisons with the PSI from *Synechocystis* sp. PCC 6803 (hereafter *Synechocystis* 6803) and *Thermosynechococcus vestitus* (formerly *Thermosynechococcus elongatus*) revealed the absence of two characteristic Chl clusters in *G. violaceus* that those authors refer to as Low1 and Low2. Note that in this work, we refer to individual Chl sites by the numbers initially assigned upon the first high resolution structure reported by Jordan et al. 2001 (*4*). Low1 corresponds to Chls A12 and A14. In *G. violaceus* PSI, A14 is absent, abolishing the Low1 low-energy Chl cluster. Low2 corresponds to Chls B31, B32, and B33. In *G. violaceus* PSI, B33 is absent, abolishing the Low2 low-energy Chl cluster. When present, these clusters cause 77 K fluorescence emission peaks at approximately 723 nm and 730 nm, whereas the 77 K fluorescence emission peak of *G. violaceus* PSI that lacks these clusters is at 695 nm. Notably, A14, which π-stacks with A12, is conserved across most oxyphototrophs, except for *G. violaceus*, suggesting that the absence of A14 is a distinctive feature of early-branching cyanobacteria. Interestingly, there is a large spectral gap between 695 and 723 nm, which raises the question of whether another PSI with an intermediate emission between these two wavelengths exists.

*Anthocerotibacter panamensis* is a representative thylakoid-free cyanobacterium that diverged from the other thylakoid-free cyanobacterial clade (*Gloeobacter* spp.) ∼1.4 billion years ago (*13, 14*). *A. panamensis* is a member of the same order, Gloeobacterales, as *Gloeobacter* spp. It was isolated as part of the microbiome of the hornworts *Anthoceros* and appeared to be a close relative of the recently described *Candidatus* Cyanoaurora vandensis, known from the metagenome of a microbial mat at the bottom of a permanently ice-covered lake in Antarctica (*15*). However, *A. panamensis* is the sole isolated and cultured species of its class, which encourages further characterization (*13, 14*). Photosynthesis in *A. panamensis* remains largely uncharacterized relative to the better-studied *G. violaceus* (*12, 16–22*). Our previous work on *A. panamensis* showed that its phycobilisome (PBS) possesses a distinctive paddle shape and preserves relict features (*23*).

Here, we perform a thorough characterization of PSI from *A. panamensis* and revisit the evolution of PSI subunits in cyanobacteria to extract novel insights into the diversification of oxygenic photosynthesis. Because the PSI of *A. panamensis* has not been studied previously, we isolated and characterized it using spectroscopic, proteomic, and phylogenetic approaches, and compared it with the PSI isolated from *G. violaceus* and the thylakoid-containing cyanobacterial model strain *Synechocystis* 6803. Through single-particle cryo-EM, we resolved the structure, identified unannotated or misannotated subunits in the previous study (*13*), and showed that PSI subunits in *A. panamensis* have followed a similar evolutionary trajectory to that of other Gloeobacterales. Lastly, we suggest energetic roles for individual Chl sites based on structural comparisons and show that *A. panamensis* and other Gloeobacterales retain a PSI that might have had an oligomerization state and subunit composition identical to that found in the most recent common ancestor of extant cyanobacteria.

## Results

### Biochemical characterization of PSI in *Anthocerotibacter panamensis*

77 K fluorescence emission spectroscopy was used to analyze whole cells with an excitation wavelength at either 440 nm for Chl molecules or 580 nm for phycobiliproteins (**fig. S1**). The emission spectrum from *A. panamensis* cells obtained with 440 nm excitation showed a peak at 688 nm and a shoulder at ∼700 nm (**fig. S1A**). For comparison, *G. violaceus* cells showed a peak at 688 nm but did not show an emission shoulder around 700 nm, and *Synechocystis* 6803 cells showed a long-wavelength peak at 725 nm (**fig. S1A**). Similarly, previous studies have demonstrated that *G. violaceus* cells do not exhibit long-wavelength fluorescence emission at 77 K, and isolated *G. violaceus* PSI trimers showed a 77 K fluorescence emission wavelength around 690 nm, similar to the emission wavelength from PSII (*12, 18*). Therefore, the emission shoulder at ∼700 nm from *A. panamensis* cells likely arises from its PSI. However, under 580 nm PBS excitation, a peak at 689 nm originating from PSII was present, while the ∼700 nm emission feature of *A. panamensis* was absent (**fig. S1B**). In contrast, *Synechocystis* 6803 cells exhibited peaks at 689 nm and 723 nm from PSII and PSI, respectively (**fig. S1B**).

This observation suggests that, at least under the conditions studied, the PBS in *A. panamensis* transfers energy to PSII but not PSI.

We isolated PSI from *A. panamensis* by sucrose density gradient centrifugation (**Fig. 1A**). Detergent-solubilized membranes were loaded onto sucrose density gradients as described in **Materials and Methods** and three fractions were observed (**fig. S2A**). Based on absorption and low-temperature fluorescence spectra compared with the previous PBS study (*23*), we concluded that Fraction 1 mainly contained carotenoids and dissociated phycobiliproteins, and Fraction 2 mainly contained phycobiliproteins and PSII (**fig. S2**). The absorption spectrum of Fraction 3 showed an absorption peak at 679 nm (**Fig. 1B**), which is consistent with the PSI absorption peak in other cyanobacteria (**fig. S3A**) and was therefore assigned as such. Note that the PSI absorption from *Synechocystis* 6803 and *T. vestitus* showed absorption features above 700 nm, which are absent in *A. panamensis* and *G. violaceus* (**fig. S3A**). The 77 K fluorescence spectrum of *A. panamensis* PSI revealed a peak at 708 nm (**Fig. 1B**), which is longer than that of *G. violaceus* (695 nm) but shorter than that of *Synechocystis* 6803 (722 nm) and *T. vestitus* (730 nm) (**fig. S3B**) (*24, 25*). Analysis of pigment extracts showed that *A. panamensis* PSI contains carotenoids canthaxanthin, echinenone, and β-carotene in a 1.0:3.3:6.5 ratio based on the integration of their peaks in the pigment analysis and their extinction coefficients. Our comparison to pigment extracts from *G. violaceus* and *Synechocystis* 6803 PSI showed that all three contain β-carotene, echinenone, and Chl *a* (**fig. S4**). Only *A. panamensis* PSI contains canthaxanthin (**Fig. 1C**, **fig. S4** and **fig. S5A**). *Synechocystis* 6803 PSI additionally contains zeaxanthin (**fig. S4** and **fig. S5B**).

**Fig. 1.**
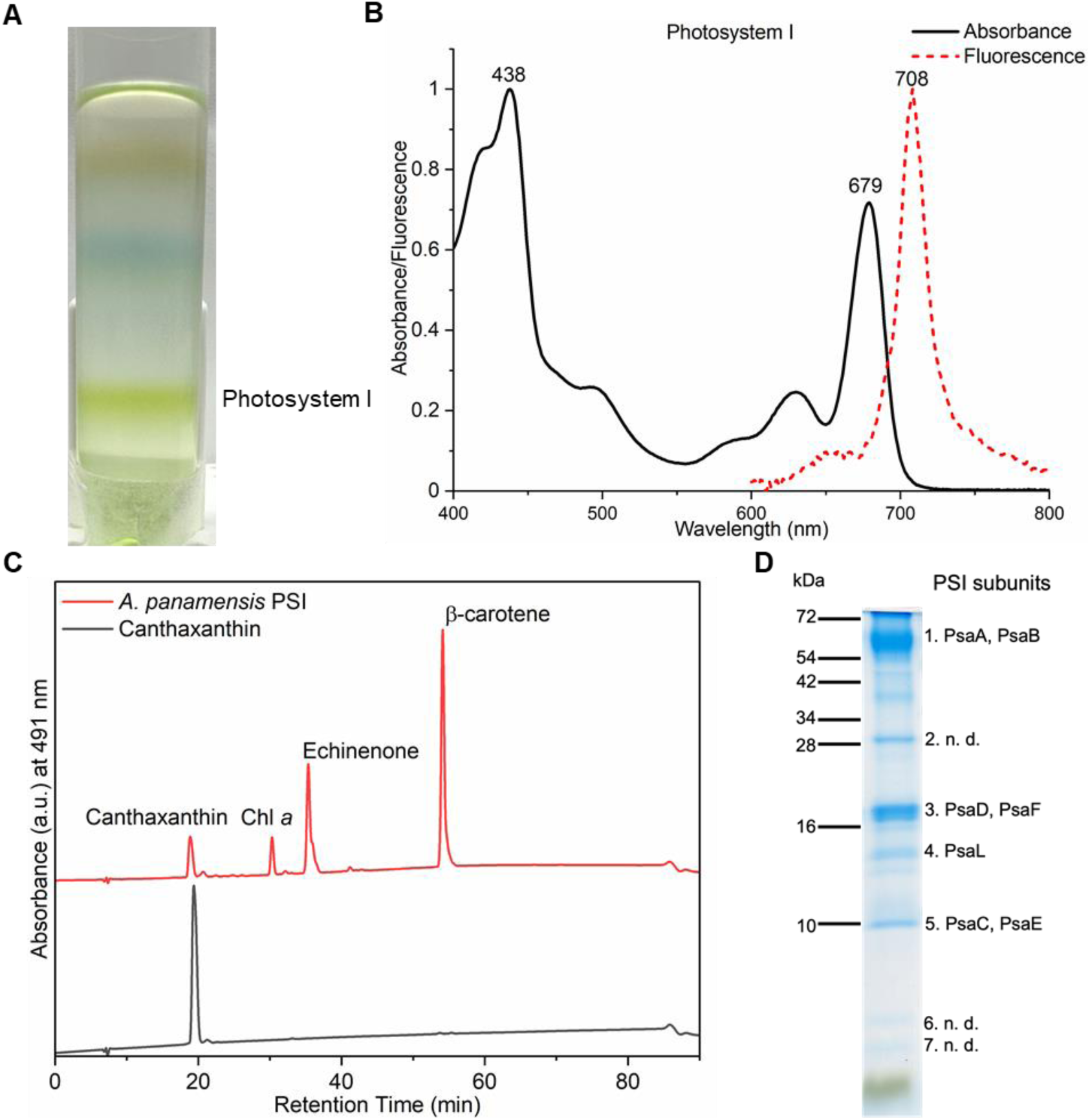
Spectral and biochemical characterization of the PSI from *A. panamensis*. (**A**) Separation of the cell membranes solubilized by *n*-dodecyl-β-D-maltoside (β-DM) by sucrose density gradient centrifugation from *A. panamensis.* The PSI fraction of *A. panamensis* is labeled. (**B**) Room-temperature absorption spectrum of *A. panamensis* PSI (solid black line). Low temperature (77 K) fluorescence emission spectrum of *A. panamensis* PSI excited at 440 nm (dashed red line). (**C**) HPLC analysis of pigments extracted from *A. panamensis* PSI. Three major pigment peaks are eluted from the PSI and are identified as canthaxanthin, Chl *a*, echinenone, and β-carotene, respectively, based on their elution time and characteristic absorption spectra (**fig. S5**). (**D**) SDS-PAGE of *A. panamensis* PSI. PSI subunits are labeled based on their molecular weights and mass spectrometry results (n. d.: PSI subunits not detected) (**table S2**).

To determine the subunit composition of *A. panamensis* PSI, we conducted in-solution liquid chromatography with tandem mass spectrometry (LC-MS/MS) on the PSI-containing Fraction 3. PsaA, PsaB, PsaC, PsaD, PsaE, PsaF, and PsaL were identified (**table S1**). Note that neither the IsiA-like PSI accessory antenna proteins nor the carotenoid-containing orange carotenoid protein were identified in the fractions. Indeed, no IsiA-like proteins are annotated in the *A. panamensis* genome or that of any other Gloeobacterales. This suggests that all the carotenoids identified in Fraction 3 are associated with PSI (*13*) (**table S1**). SDS-PAGE analysis of the *A. panamensis* PSI fraction revealed separated protein bands, in which PSI subunits PsaA, PsaB, PsaC, PsaD, PsaE, and PsaF were identified by in-gel LC-MS/MS (**Fig. 1D** and **table S2**). Based on our subunit analysis, we did not detect common PSI subunits PsaI, PsaJ, PsaK, PsaM, or PsaX. It should be noted, however, that small transmembrane subunits such as these are challenging to detect by LC-MS/MS.

### Cryo-EM structure of *A. panamensis* PSI

To reveal the molecular structure of *A. panamensis* PSI, we performed electron microscopy. Negative staining followed by transmission electron microscopy (TEM) and 2D image classification showed that the *A. panamensis* PSI in Fraction 3 exists in a trimeric oligomeric state (**fig. S6**), like many other cyanobacterial species. To obtain higher resolution structural data, we performed cryo-EM. Initial screening on a 200 kV Glacios microscope confirmed the trimeric arrangement of *A. panamensis* PSI and helped to identify optimal plunge freezing conditions (**fig. S7** and **Materials and Methods**).

High-resolution data collection was performed on a 300 kV Titan Krios (**fig. S8**) which led to the structural determination of *A. panamensis* PSI at 2.4 Å global resolution (**Fig. 2**, **fig. S9**, and **table S3**). Each PSI monomer exhibits pseudo-C2 symmetry about its central core, which is composed of subunits PsaA and PsaB. Transmembrane subunits PsaF, PsaI, PsaJ, PsaL, and PsaM, and stromal side soluble subunits PsaC, PsaD, and PsaE were also identified. Although PsaI, PsaJ, and PsaM were not detected in the LC-MS/MS analysis, PsaM was expected to be present based on its identification in the *A. panamensis* genome (*13*). The more divergent PsaI and PsaJ subunits were not identified in the genome until recently (*26*), which is consistent with their clear identification in the cryo-EM map (**Fig. 2D** and **Fig. 2E**). The *psaI* subunit for *A. panamensis* and its close relative, *Candidatus* C. vandensis, were found downstream from a gene annotated as “response regulator transcription factor” (WP_287128339.1 and WP_218081160.1, respectively) and the gene *psaJ* was found adjacent to *psaF* in both strains. Thus, compared to the subunit composition of other PSI structures, an important characteristic of *A. panamensis* PSI is that it lacks subunits PsaK and PsaX, the common locations of which are shown in **Fig. 2A**. This is also the case for *G. violaceus* PSI, and therefore may be a common trait among PSI from thylakoid-free cyanobacteria.

**Fig. 2.**
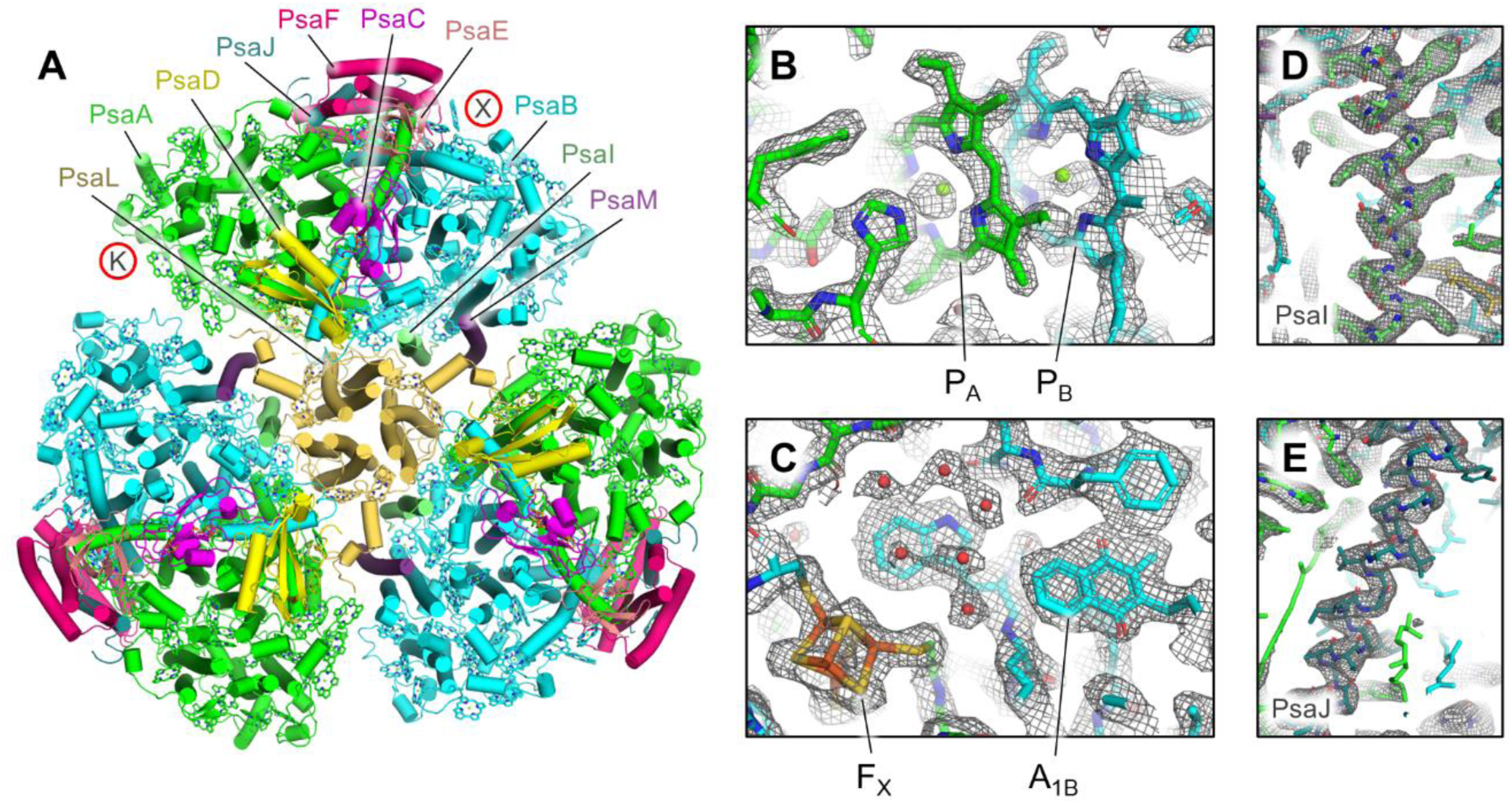
Cryo-EM structure of *A. panamensis* PSI. (**A**) Structure of the trimeric *A. panamensis* PSI complex viewed from the stromal side. Cofactors except Chls are hidden for clarity, and those are shown as tetrapyrrole rings only. Note that the PsaK and PsaX subunits found in PSI complexes from some other cyanobacterial species are not present in *A. panamensis*. Their locations relative to other PSI structures are designated by the letters K and X in red circles. (**B**) Model within the sharpened map showing the vicinity of P700 which is composed of the PA (a Chl *a’* molecule) and PB (a Chl *a* molecule). (**C**) Model within the sharpened map showing the vicinity of FX (a [4Fe-4S] cluster), A1B (a menaquinone-4 molecule), and nearby waters (red spheres). (**D**) Model within the unsharpened map of PsaI that was previously suggested not to be found in *A. panamensis*. (**E**) Model within the unsharpened map of PsaJ that was previously suggested not to be found in *A. panamensis*.

The cofactor composition based on the structural analysis of *A. panamensis* PSI is similar to other cyanobacterial PSI complexes, where each PSI monomer binds 88 Chl *a*, 1 Chl *a’*, 2 quinones, 22 carotenoids, three [4Fe-4S] clusters, and numerous lipids (**table S4**). *G. violaceus* PSI contains menaquinone-4 (MQ-4) in its quinone-binding sites instead of phylloquinone-4 (PhQ-4) found in thylakoid-containing cyanobacteria such as *Synechocystis* 6803 and *T. vestitus*. Structures of these two quinones are shown in **fig. S10**. To determine the quinone type found in *A panamensis* PSI, we first compared the quinone tail orientations of *A. panamensis* PSI with PSI structures from *G. violaceus* and three thylakoid-containing cyanobacteria (**Fig. 3A**). The quinone tails found in *A. panamensis* PSI most closely match those found in *G. violaceus* PSI, suggesting that they are MQ-4 molecules. To further investigate this possibility, we separated cofactors from *A. panamensis* PSI (possibly containing MQ-4) and *Synechocystis* 6803 PSI (known to contain PhQ-4 and examined their absorption spectra (**Fig. 3B** and **Fig. 3C**). The characteristic quinone absorbance spectrum and elution time from *A. panamensis* PSI are consistent with a MQ-4 standard, whereas the characteristic quinone absorbance spectrum and elution time from *Synechocystis* 6803 are consistent with a PhQ-4 standard. This observation provides strong support that, like *G. violaceus* PSI, *A. panamensis* PSI contains MQ-4 in its quinone-binding sites, and that this is more broadly a feature of thylakoid-free cyanobacteria.

**Fig. 3.**
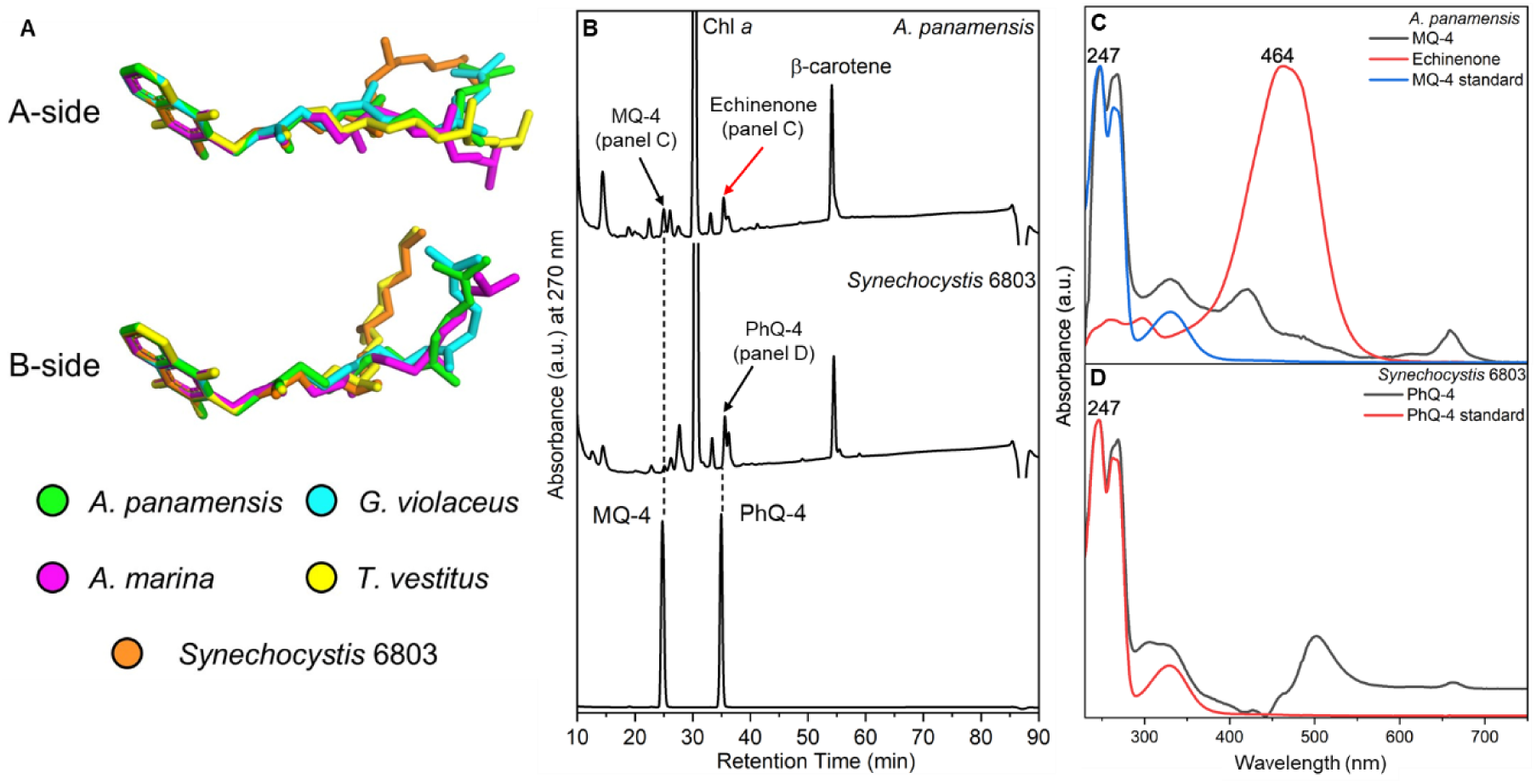
The structure and HPLC analysis of MQ-4 and PhQ-4 in the PSI. (**A**) Structures of quinones from selected PSI structures. C atoms in the quinone rings of the quinones in PSI structures from *A. panamensis*, *G. violaceus*, *A. marina*, *T. vestitus*, and *Synechocystis* 6803 are superimposed. The latter four correspond to PDBs 7F4V, 7COY, 1JB0, and 5OY0, respectively. (**B**) HPLC elution profile of extracts from PSI complexes of *A. panamensis* and *Synechocystis* 6803 (top two traces, respectively), and a standard containing MQ-4 and PhQ-4 (bottom trace). The black arrow indicates the peak of quinones in the PSI extracts. The red arrow indicates the peak of echinenone in the *A. panamensis* PSI extract. (**C**) Absorption spectrum of MQ-4 standard, MQ-4 and echinenone from PSI of *A. panamensis*. (**D**) Absorption spectrum of PhQ-4 standard and PhQ-4 from PSI of *Synechocystis* 6803.

Based on the ratios of canthaxanthin, echinenone, and β-carotene determined from pigment analysis, we expect ∼2, 7, and 13 of those carotenoids in each PSI monomer, respectively. In *A. panamensis* PSI, consistent with other PSI structures, the carotenoids are highly flexible and exhibit relatively weak cryo-EM density. Therefore, distinguishing between the three carotenoids is challenging due to their minor structural differences (**fig. S11**). Consequently, due to the insufficiency of the cryo-EM density for definitive carotenoid type assignment, we modeled all but one carotenoid as β-carotene, the most prominent carotenoid identified in pigment analysis (**Fig. 1C)**. However, the ring headgroup of one carotenoid located near the monomer-monomer interface (**Fig. 4A**) was relatively well defined, and appeared to exhibit signal greater than would be expected for a CH2 group at the C4 position of the ring closest to a nearby Chl site (**Fig. 4B**). This suggests that it is a keto-carotenoid, either canthaxanthin or echinenone. To investigate this possibility further, we performed a more quantitative analysis of the cryo-EM map (*27*) (**Fig. 4**): we sampled signal amplitude in the map at increments moving away from ring carbon positions C2, C3, C4, and C5. The C2, C3, and C5 positions are CH2, CH2, and C-CH3 in each of the possible carotenoid types (canthaxanthin, echinenone, and β-carotene) (**Fig. 4C**). As expected, the C2 (CH2) and C3 (CH2) signal amplitude drops off steeply compared to C5 (C-CH3). β-carotene has CH2 at position C4 on both rings, echinenone has CH2 at C4 on one ring and C=O on the other, and canthaxanthin has C=O at position C4 on both rings. Rather than the C4 signal dropping off steeply like positions C2 and C3 known to be CH2, it instead drops off slowly, but faster than C5 known to be C-CH3. In this context, the C4 profile is more consistent with C=O (canthaxanthin and echinenone) than CH2 (β-carotene). Thus, this carotenoid was tentatively modeled as the keto-carotenoid with the highest concentration, echinenone, although it is possible that it could instead be a canthaxanthin molecule.

**Fig. 4.**
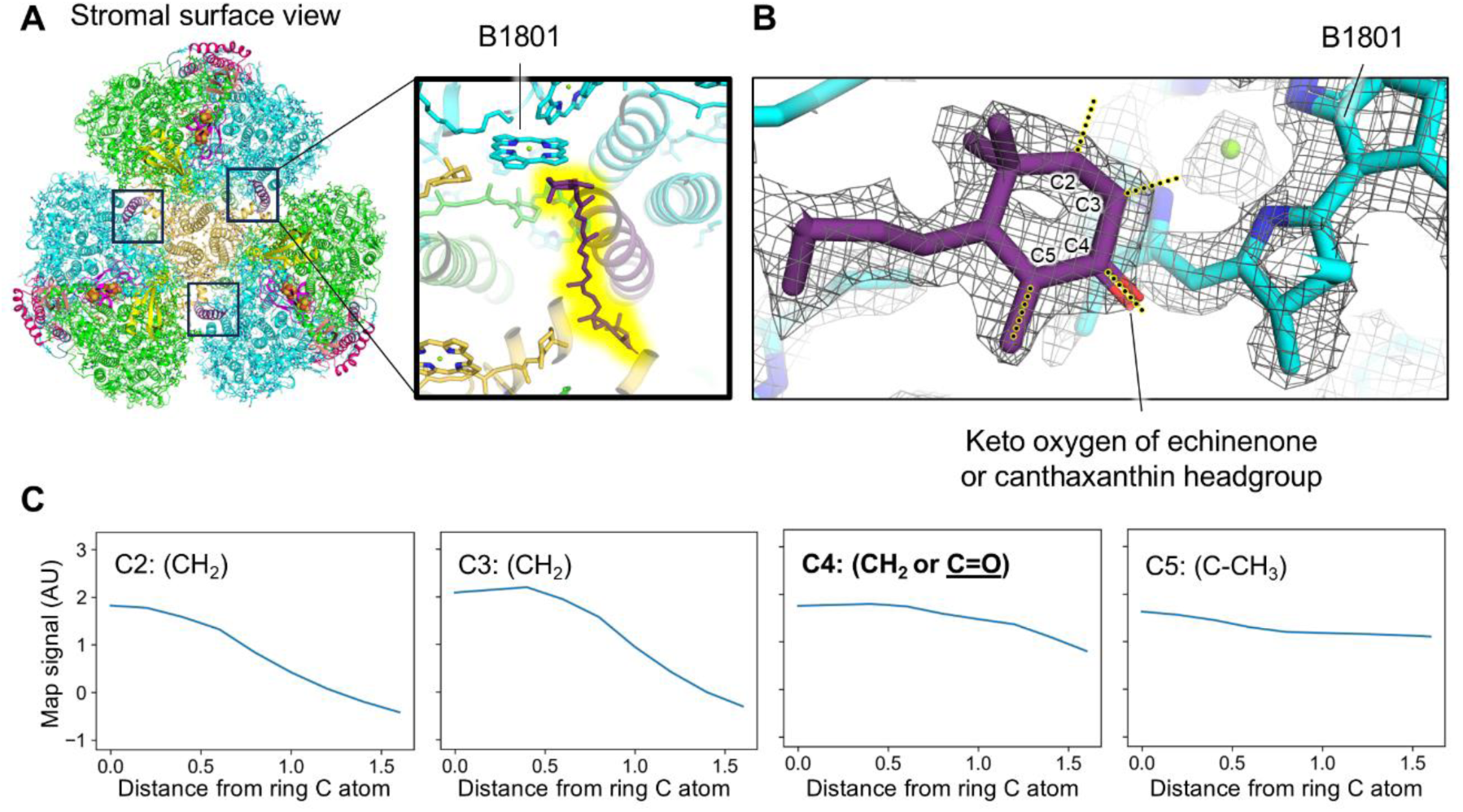
Location of the echinenone or canthaxanthin molecule in the *A. panamensis* PSI structure. (**A**) *A. panamensis* PSI trimer shown from a stromal view. Boxes near the monomer-monomer interfaces designate the echinenone or canthaxanthin locations. The top right box corresponds to the magnification. In the magnification, the echinenone or canthaxanthin is highlighted in yellow and the nearby Chl B1801 is labeled. (**B**) Model within the sharpened map of the echinenone or canthaxanthin headgroup nearby Chl B1801. (**C**) Scans of cryo-EM map signal corresponding to the yellow highlighted dotted lines in panel B. The X-axis is reported in units of Å.

### Structural comparison of *A. panamensis* PSI with other PSI structures

We had previously shown that PsaI and PsaJ are mostly unannotated in the genomes of *G. violaceus*, *G. morelensis*, *G. kilaueensis*, *A. panamensis*, and the metagenome assembled genome (MAG) of *Candidatus* C. vandensis. Some of these were previously reported in Gisriel et al. 2023 (*26*), except for PsaI in *Candidatus* C. vandensis and PsaJ in *A. panamensis*. Here, we completed the set for both.

To compare *A. panamensis* PSI subunits with PSI from other cyanobacteria whose structures are known, we calculated the sequence identities and root-mean square deviation (RMSD) of individual PSI subunits from *G. violaceus*, *T. vestitus*, and *Synechocystis* 6803 (**table S5**). In nearly all cases, *A. panamensis* PSI subunits were more similar to *G. violaceus* than they were to *T. vestitus* or *Synechocystis* 6803. For a more in-depth comparison of *A. panamensis* PSI subunits to those from *G. violaceus*, we calculated the ratio of sequence identity (larger=more similar) to RMSD (lower=more similar) and plotted the data from highest to lowest (**fig. S12**). The most similar subunits are PsaC and PsaD found on the stromal side of the complex to which electron acceptors bind. The least similar subunits are transmembrane subunits PsaL, PsaF, and PsaJ (**fig. S12** and **fig. S13**). These subunits may be under less selective pressure to maintain certain residues due to their longer distance away from the electron transfer chain cofactors. We also calculated the surface electrostatics maps for PSI structures from *A. panamensis*, *G. violaceus*, *T. vestitus*, and *Synechocystis* 6803 (**fig. S14**). All are relatively similar, which probably relates to the ubiquitous need for binding of soluble electron donors and acceptors.

Although many Chl sites are known to be conserved in PSI from different species, the variability in 77 K fluorescence maxima from PSI among cyanobacterial strains (**fig. S3B**) suggests that some Chl sites may not be conserved and/or that some Chls (or groups of Chls) exhibit variability in their site energies. The availability of various PSI structures allows for comparisons of Chl sites that give rise to such spectral features. Indeed, this opportunity was leveraged when the structure of PSI from *G. violaceus* was determined (*12*), which lack low-energy Chls. As described above, it was suggested that two sites, called Low1 and Low2, were absent in *G. violaceus* compared to some thylakoid-containing cyanobacterial PSI species (*12*).

The acquisition of the *A. panamensis* PSI 77 K emission spectrum (**Fig. 1C** and **fig. S3B**), whose peak maximum is red-shifted compared to *G. violaceus* PSI, but blue-shifted compared to PSI from thylakoid-containing cyanobacteria *T. vestitus* and *Synechocystis* 6803, and the cryo-EM structure allow for a further analysis of the low-energy Chls in PSI. We compared the Chl sites of *A. panamensis* PSI to PSI from *G. violaceus*, *Synechocystis* 6803, and *T. vestitus* (**Fig. 5**). Two Chl sites are present in *A. panamensis* PSI that are absent in *G. violaceus* PSI: A14 and J3 (**Fig. 5**, left panels). First, this suggests that one or both of the Chls in site A14 or J3 of *A. panamensis* PSI red-shifts its 77 K fluorescence spectrum. Chl A14, along with Chl A12, is part of the two-Chl cluster termed “Low1” that was recently suggested to be a low-energy Chl site present in *Synechocystis* 6803 and *T. vestitus* (*12*). Thus, the presence of Chl A14 in *A. panamensis* PSI likely contributes to the red shift in the 77 K fluorescence maximum relative to *G. violaceus*, supporting the hypothesis that “Low1” is indeed a red Chl site.

**Fig. 5.**
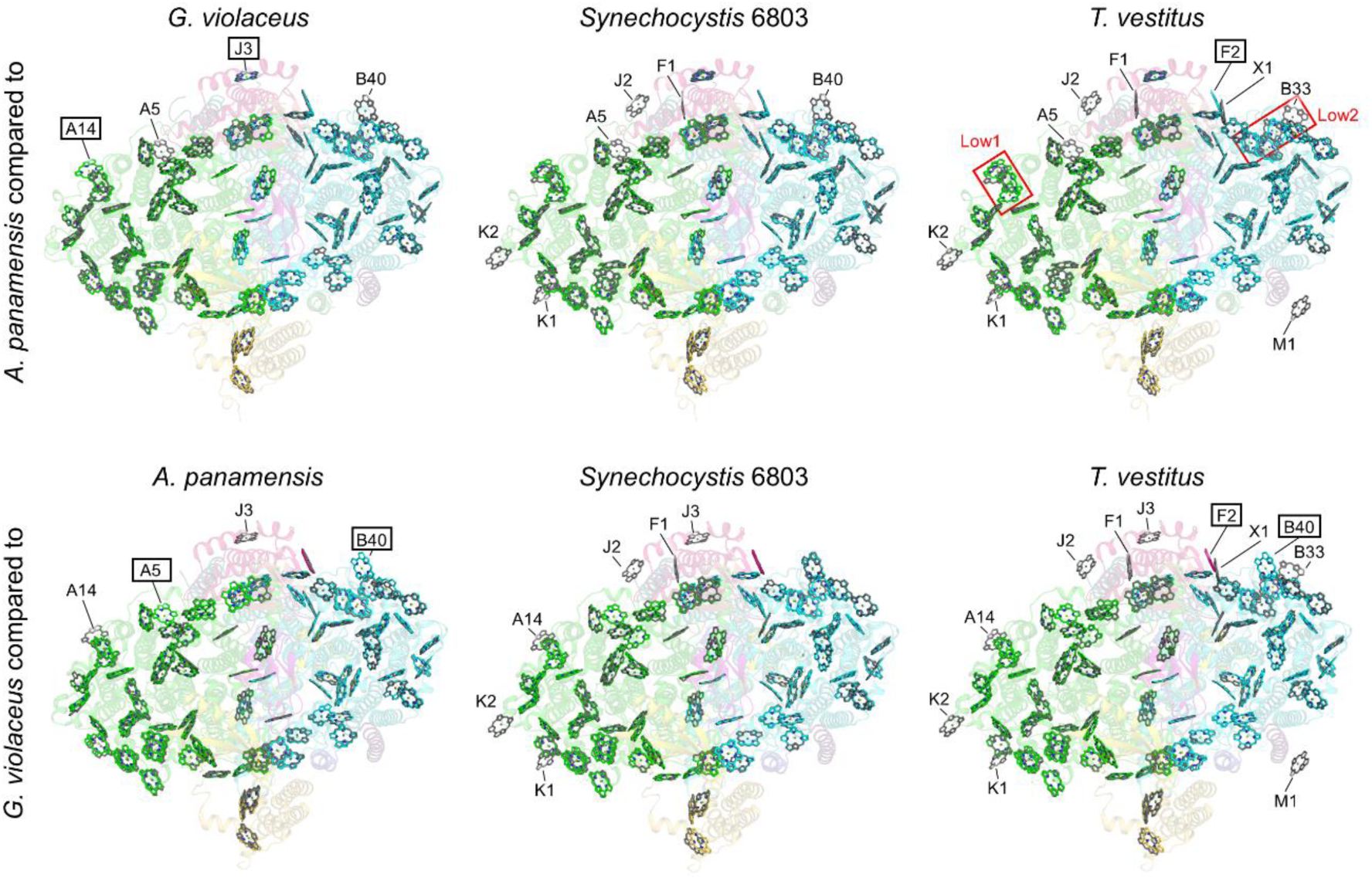
Comparison of Chl sites among selected cyanobacterial species. Each panel is a superposition of Chl sites in two structures. The top row shows comparisons of the *A. panamensis* Chl sites (colored) compared to Chl sites from *G. violaceus*, *Synechocystis* 6803, and *T. vestitus* (grey). The bottom row shows comparisons of the *G. violaceus* Chl sites (colored) compared to Chl sites from *A. panamensis*, *Synechocystis* 6803, and *T. vestitus* (grey). Labeled sites are different between the two structures. For the top row, those labels with boxes are present in *A. panamensis*, but not in the other structure. Those labels without boxes are absent in *A. panamensis*, but present in the other structures. For example, in the comparison of *A. panamensis* PSI to *G. violaceus* PSI in the top left panel, sites A14 and J3 are found in *A. panamensis*, but not in *G. violaceus*. Sites A7 and B40 are found in *G. violaceus*, but not in *A. panamensis*. For the bottom row, the same rules apply, but for *G. violaceus* instead of *A. panamensis*. For example, in the bottom middle panel *Synechocystis* 6803 PSI contains all the sites found in *G. violaceus* PSI, and additionally K1, K2, A14, J2, F1, and J3 that are not found in *G. violaceus*, so none of the labels are boxed.

Correspondingly, the protein sequence of PsaA nearby this site is generally conserved in *A. panamensis*, *Synechocystis* 6803, and *T. vestitus*, but not in *G. violaceus* where Chl A14 is absent (**fig. S15**). The Chl in site J3 is also present in *A. panamensis*, *Synechocystis* 6803, and *T. vestitus* PSI, but absent in *G. violaceus* PSI, so it may also contribute to red-shifting. Second, there are two Chl sites absent in *A. panamensis* PSI but present in *G. violaceus* PSI: A5 and B40 (**Fig. 5**). This observation suggests that neither of these Chls contribute substantially to red shifting of the 77 K fluorescence maximum. For B40, this is consistent with it also not being present in *T. vestitus* PSI, which has the most red-shifted 77 K fluorescence maximum of the strains we compared here.

Whereas *A. panamensis* and *G. violaceus* PSI both have only 89 Chls per monomer, *Synechocystis* 6803 and *T. vestitus* PSI have 95 and 96 Chls per monomer, respectively, and their 77 K fluorescence maxima are longer than PSI from *A. panamensis* and *G. violaceus*. This suggests that at least some of the additional Chls bound in PSI from these thylakoid-containing cyanobacteria contribute to red-shifting of their 77 K fluorescence maxima. Possible candidates are Chls K1, K2, J2, and F1, all of which are present in PSI from the thylakoid-containing cyanobacterial species but not in PSI from *A. panamensis* or *G. violaceus*. Furthermore, the Chls in sites X1, B33 (previously suggested to be in “Low2” (*12*), and M1 are all found only in *T. vestitus*, so they are also candidates for low energy Chl sites. Finally, it is noteworthy that Chl A5 is present in PSI from the thylakoid-containing cyanobacterial species but also in *G. violaceus* PSI, and Chl F2 is present in both *A panamensis* and *G. violaceus* PSI, so both probably do not contribute to red-shifting of the 77 K fluorescence maximum.

### Phylogenetic analysis shows conservative evolution in *A. panamensis*

To understand the position of *A. panamensis* PSI within the context of the evolution of cyanobacteria, we performed phylogenetic analyses of each subunit found in the complex, as well as PsaK and PsaX. We compiled an updated sequence dataset extracted from over 9 million cyanobacterial proteins in genome and metagenome assemblies currently at the National Center for Biotechnology Information (NCBI) (*28*). Our dataset included sequences from seven strains classified as Gloeobacterales, including *A. panamensis*, *G. violaceus*, *Gloeobacter morelensis*, *Gloeobacter kilaueensis*, *Candidatus* C. vandensis (the closest relative of *A. panamensis*) and two unclassified metagenome assembled genomes (ES-bin-313 and ES-bin-141) from two relatives of the well-known *Gloeobacter* spp. retrieved from Greenland (*29*). All PSI subunits in the Gloeobacterales showed strong affiliation with each other (**fig. S16**) regardless of variation in sequence length and rates of evolution. We found no evidence for duplication of existing PSI subunits or gain of known PSI subunits from other cyanobacteria via horizontal gene transfer.

Gloeobacterales are considered to retain a greater number of ancestral traits than other cyanobacteria. For this to be true, it is required that, on average, Gloeobacterales have evolved at a comparatively slower rate than other cyanobacteria. Given that PsaA and PsaB originated from a gene duplication event, the level of sequence identity of PsaA compared with PsaB must decrease with time as amino acid substitutions accumulate in each subunit through cyanobacteria diversification. Therefore, it can be hypothesized that if Gloeobacterales have evolved slower than other cyanobacteria, the level of sequence identity of PsaA v PsaB should be greater in Gloeobacterales—because they have accumulated less change—than in thylakoid-containing cyanobacteria. To test this hypothesis, we calculated ancestral sequences for the ancestors of PsaA and PsaB (marked with green spheres in **Fig. 6**) across well conserved regions (676 positions) in an alignment of 1,074 PsaA and PsaB sequences. We found that PsaA and PsaB in the most recent common ancestor (MRCA) of extant cyanobacteria shared about 55% sequence identity. Today PsaA v PsaB share 50% sequence identity in *G. violaceus*, 48% in *A. panamensis* and *T. vestitus*, 46% in *Synechocystis* 6803, and 43% in the faster-evolving *Prochlorococcus marinus* (see **Table 1**). This suggests that if PsaA and PsaB in Gloeobacterales evolved slower than in other cyanobacteria, the difference is only minor.

**Fig. 6.**
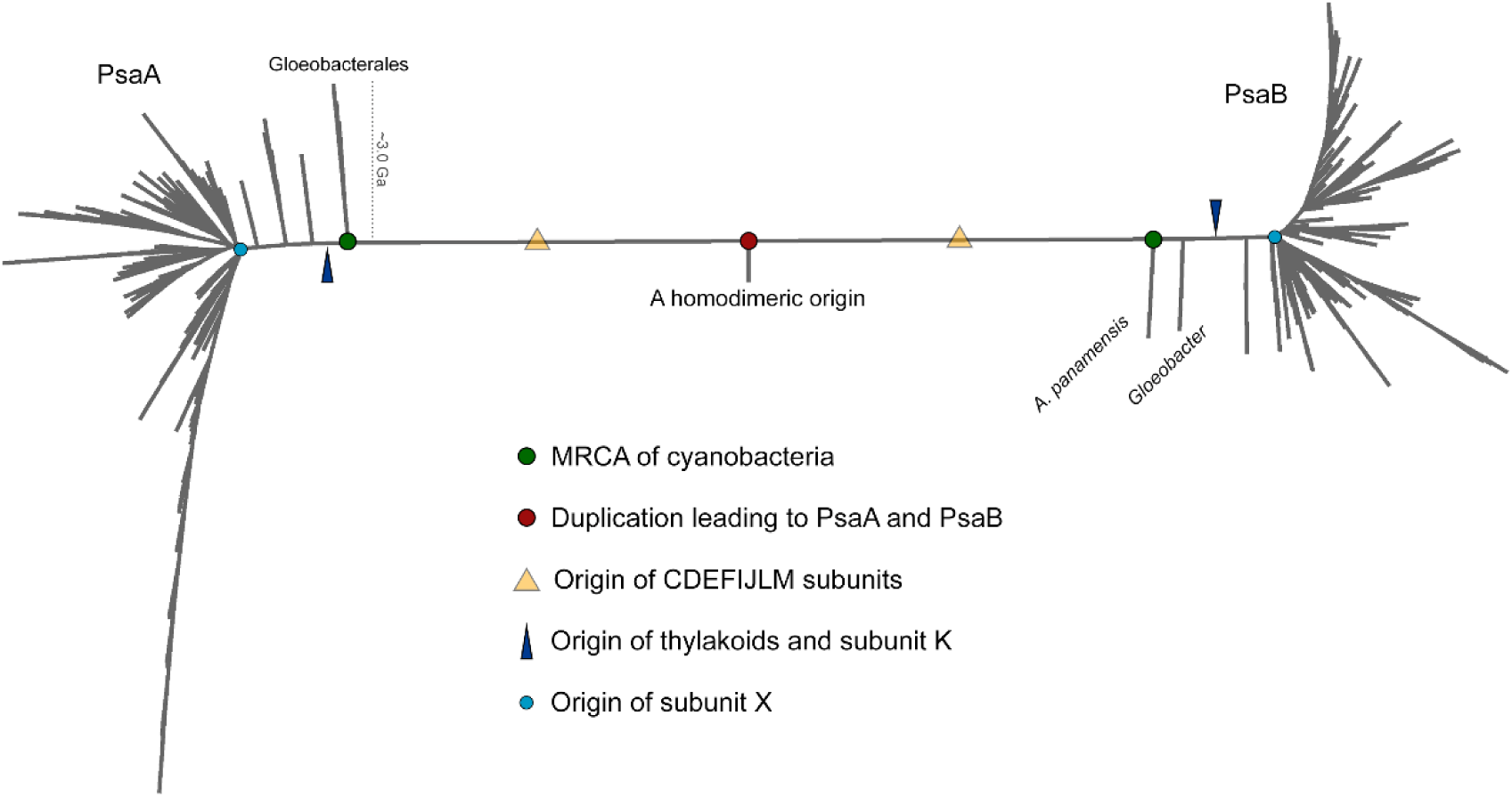
Phylogenetic tree of PsaA and PsaB. Some of the key events in the evolutionary history of PSI are overlaid with PsaA and PsaB evolution. PsaA and PsaB originated from an ancient gene duplication when the photosystem was homodimeric. The most recent common ancestor (MRCA) of Cyanobacteria, represented by green circles, often thought to have originated before the Great Oxidation Event, inherited both PsaA and PsaB and already had heterodimeric PSI. All cyanobacteria retain subunits PsaC, PsaD, PsaE, PsaF, PsaI, PsaJ, PsaL, and PsaM, which implies that these were added to PSI before the MRCA of cyanobacteria and a capacity for trimerization (yellow triangle). On PsaA, Gloeobacterales, including *A. panamensis*, its close relative *Candidatus* C. vandensis, and species of *Gloeobacter* make the earliest branching clade. On PsaB, Gloeobacterales is separated into two distinct clades, a phenomenon that had been observed before (*13*), but likely represents a long branch attraction artifact triggered by the long branch that separates PsaA and PsaB. PsaK appears to have originated after the branching event leading to the Gloeobacterales (dark blue triangle), while PsaX appears to have originated close to or during the major cyanobacteria radiation leading to microcyanobacteria and macrocyanobacteria (light blue circle), as defined by Sanchez-Baracaldo et al. (*1*).

**Table 1.**
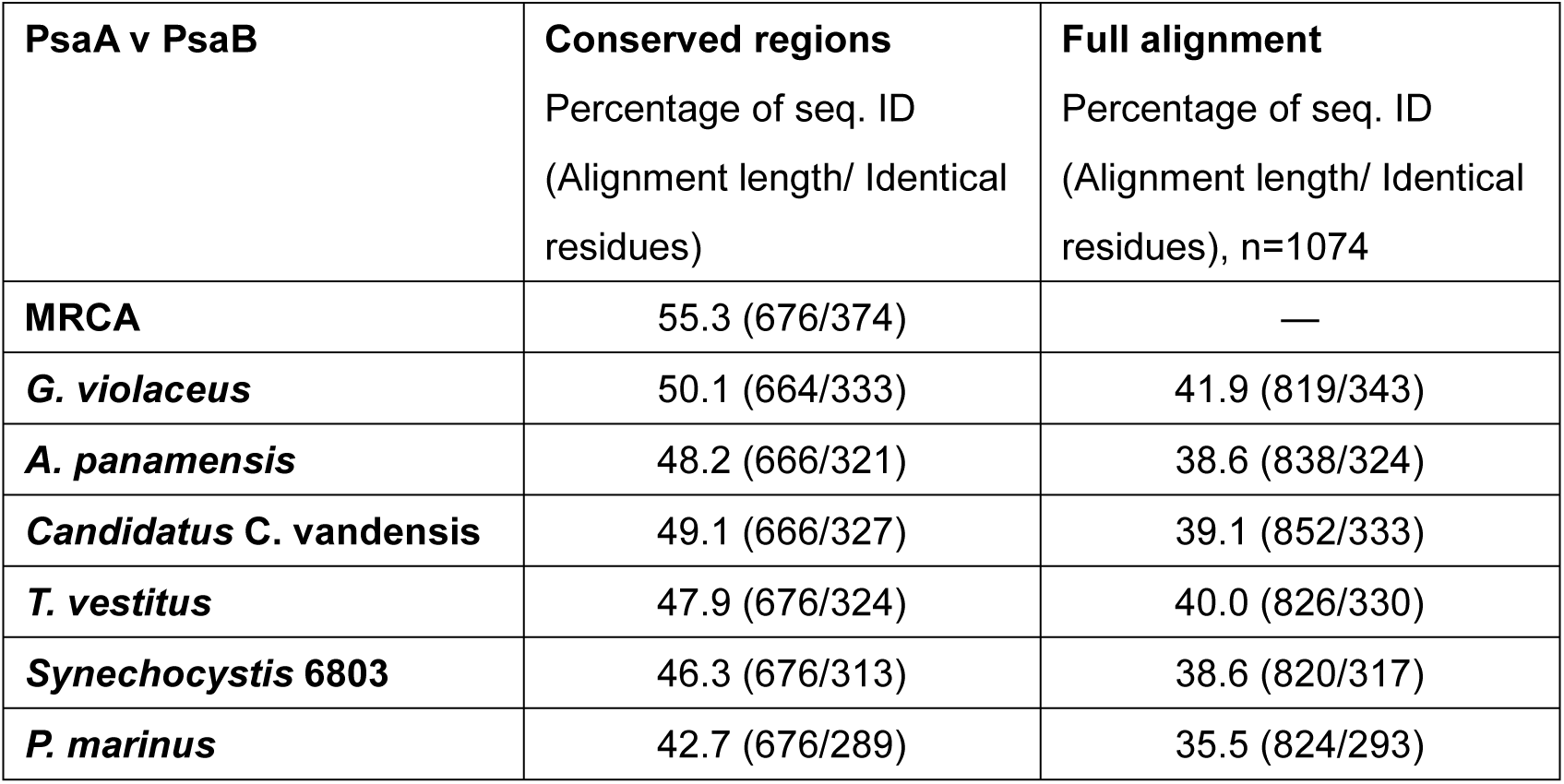
Percentage of sequence identity between PsaA and PsaB.

The phylogeny of a tree combining both PsaA and PsaB placed the Gloeobacterales as the most basal branches as expected. However, in the PsaB side, the Gloeobacterales was split into separate clades having *A. panamensis* and *Candidatus* C. vandensis as the earliest branching clade followed by *Gloeobacter* spp. (**Fig. 6**). This phenomenon had been reported before and likely represents a long-branch attraction artifact (*15*), rather than a true signal, as the effect disappears when each tree is built separately.

We also observed that PsaC is the most highly conserved subunit of PSI (**fig. S16**), consistent with our structure-based comparison (**fig. S12A**), with only three substitutions in 81 positions when comparing *G. violaceus* with *A. panamensis*, despite separating ca. 1.4 billion years ago. Only five substitutions are noted between *Synechocystis* 6803 and *A. panamensis* even though their MRCA could comfortably be 3.0 billion years ago (*30, 31*). These findings suggest a rate of amino acid change in PsaC of ∼2 residue substitutions per billion years. It is worth noting that the PsaC sequence from the FaRLiP- capable *Halomicronema hongdecloris* clustered in the phylogeny of PsaC next to those of other Gloeobacterales, featuring only 3 amino acid substitutions when compared with *A. panamensis*, with no branch length, suggesting that the PsaC sequence in the ancestor of Gloeobacterales may be identical to that of the MRCA of extant cyanobacteria.

Another notable finding relates to the phylogeny of PsaL (**fig. S17**), which enables the oligomerization of PSI complexes. In some cyanobacterial genomes an isoform of PsaL is found, which is fused with an IsiA-like Chl-binding protein. It was shown recently that *Nostoc* sp. PCC 7120 expresses this IsiA-PsaL fusion under iron-deficient conditions, binding to PSI monomers and disrupting oligomer formation (*32*). We found this fused PsaL to cluster early within cyanobacteria evolution after the divergence of Gloeobacterales and the early-branching *Synechococcus*, suggesting that its evolution may have been facilitated by the origin of thylakoid membranes. Furthermore, we examined the phylogeny and distribution of PsaK, which is also absent in Gloeobacterales but has otherwise a broad distribution, including most other basal clades (**fig. S17**). The distribution of PsaX is more limited than that of PsaK. It is absent from all basal clades (Gloeobacterales, the *early-branching Synechococcus*, Pseudanabaenales, Gloeomargaritales) except for some strains of *Thermosynechococcus*. PsaX appears to be mostly found in heterocystous cyanobacteria and their relatives, as well as several other macrocyanobacteria clades.

## Discussion

The analyses performed herein provide numerous unique insights into the diversity and evolution of photosynthesis. It was recently suggested that, based on the structure of *G. violaceus* PSI, the absence of “Low1” is a characteristic of primordial cyanobacteria (*12*). Our structural and phylogenetic analyses do not support this hypothesis because the residues that bind Chl A14 nearby Chl A12 are conserved in the ancestral sequence reconstruction (**fig. S15**). This highlights an important point: one should be cautious when referring to thylakoid-free cyanobacteria as “primitive”, as all extant cyanobacteria are equally distant in time from the MRCA of cyanobacteria (**Fig. 6**). The differentiating factor is the rate of evolution, and our data suggest only minor differences in the rates of evolution for Gloeobacterales compared to other cyanobacteria. Furthermore, in some respects, Gloeobacterales have diversified more than other cyanobacteria. A prime example is that although the *A. panamensis* phycobilisome retains ancestral phycobilisome characteristics such as the absence of phycocyanin rods, ApcD, and ApcF, it also exhibits novel phycocyanin chains and a heptacylindrical core (*23*).

The fluorescence peak maxima among PSI from various cyanobacterial species exemplifies the diversity of light-harvesting strategies. Previous studies have shown that overall trapping kinetics decrease with increasing low-energy Chls in PSI complexes (*8, 33, 34*). Thus, although low-energy Chls in PSI can expand the light-harvesting cross-section to longer wavelengths, they also inevitably decrease efficiency (*34, 35*), the balance of which is likely driven by ecological resources and perhaps even cellular architecture. The available membrane space for the photosynthetic apparatus in the thylakoid-free Gloeobacteriales is restricted to the plasma membrane (*13, 23, 36*). To overcome this limitation, these cyanobacteria adapted to produce large phycobilisomes to harvest more light for driving PSII function (*23*). Decreasing the number of low-energy Chls in PSI might have been driven by the need for increased PSI trapping kinetics.

Nevertheless, this adaptation also results in the loss of photoprotection capabilities, which is likely why Gloeobacteriales can only be cultivated in low-light environments (*13, 36–38*). In contrast to *G. violaceus*, *A. panamensis* can tolerate a somewhat higher light intensity of up to 100 μmol photons m^−2^ s^−1^ (*13*), which may be attributed to the additional low-energy Chl site in *A. panamensis* PSI. (**Fig. 5**, **fig. S3**, and **fig. S15**).

Recent phylogenomic studies have suggested that, unlike known Gloeobacterales, the MRCA of cyanobacteria likely inhabited shallow marine habitats that were probably well illuminated environments (*31*), suggesting that PSI adaptations to high or low-light intensity environments may be species-dependent and is likely have occurred repeatedly and in either direction. An evolutionary characteristic that is seen in PSII (*39*), where tuning of the photosystem energetics to either condition can be achieved with relatively few amino acid substitutions such as a change of Glu for Gln, or vice versa, at position 130 of the D1 subunit.

Another key finding of this work is that *A. panamensis* PSI contains MQ-4 at its quinone-binding sites, a feature that it shares with *G. violaceus* PSI. Therefore, this may be a common trait among thylakoid-free cyanobacteria. However, MQ-4 binding in the PSI electron transfer chain is also observed in the cyanobacterium *Synechococcus sp.* PCC 7002 (*40, 41*), and the red alga *Cyanidium caldarium* (*42*), suggesting that MQ-4 is more widespread among oxygenic phototrophs than is currently recognized. MQ-4 and PhQ-4 share similar biosynthetic pathways, both utilizing 1,4-naphthoquinone as a precursor (*41, 43*). However, they differ structurally in the saturation level of the aliphatic side chain attached at the 3-position (**fig. S10**). If the presence of MQ-4 in PSI was a characteristic of the MRCA of cyanobacteria, it is possible that PhQ-4 biosynthesis developed in many thylakoid-containing cyanobacteria over time, which may confer their advantage to tolerate high light, as mutants that lack PhQ-4 in *Synechocystis* 6803 are unable to grow in high light conditions (*41*).

Another interesting observation is the unique carotenoid composition of *A. panamensis* PSI, containing β-carotene, echinenone, and canthaxanthin. This is unlike other cyanobacteria that typically contain the former two but not the latter (*44, 45*). As far as we know, the trimeric PSI that contains canthaxanthin or echinenone is only found *A. panamensis*. The presence of canthaxanthin may reflect a specialized light-harvesting or photoprotective adaptation in *A. panamensis*. In studies of tetrameric PSI structures, carotenoids such as myxoxanthophyll, echinenone, and canthaxanthin are enriched at oligomeric interfaces, particularly under high-light conditions, where they play crucial roles in light harvesting and photoprotection (*46*). Other research has shown that in the trimeric PSI of *Synechocystis* 6803, zeaxanthin and echinenone were detected by HPLC, and mutants lacking these carotenoids have reduced PSI stability (*45, 47*). Canthaxanthin or echinenone was identified at the monomer-monomer interface (**Fig. 4**) in the *A. panamensis* PSI structure, suggesting similar roles of this molecule in photoprotection and/or structural stabilization. Additionally, carotenoids located at different sites within the PSI complex may have distinct functions—either dissipating excess energy or aiding in light harvesting, depending on light conditions (*48, 49*). The diverse carotenoid composition in *A. panamensis* likely allows it to adapt to varying light intensities, a particularly important trait given its known high-light tolerance (*13*). Future studies should investigate the dual roles of carotenoids in *A. panamensis* PSI and their broader implications for carotenoid diversity and function in cyanobacteria.

Our evolutionary analysis suggests that, while the rates of PSI evolution in Gloeobacterales do not stand out as particularly slow relative to other cyanobacteria, there is indeed little innovation in terms of PSI architecture when *A. panamensis* is compared with *G. violaceus* and with other trimers. There is neither evidence for the emergence of PSI subunit isoforms that could offer new capabilities, unlike D1 in PSII (*38, 50*), nor is there evidence for the recruitment of novel subunits into PSI. Remarkably, the genome of *G. violaceus* appears to encode a PsaB subunit fused with an outer membrane protein, presenting a unique C-terminal 155 amino acid extension (*20*), which could have represented a novel feature. The translation of this extension was apparently confirmed via mass spectrometry in purified PSI and it was hypothesized that it could anchor PSI to the peptidoglycan layer (*20*), but this extension was not found in the respective structure (*12*). Therefore, although sequence change has occurred over the past few billion years, it can be concluded that at an architectural level, the MRCA of cyanobacteria had heterodimeric PSI cores that assembled into trimers and had a subunit composition similar to those found in *A. panamensis* and *G. violaceus*, lacking PsaK and PsaX. After the MRCA of cyanobacteria began to diversify into the extant clades, within the lineage leading to Gloeobacterales, relatively few architectural changes occurred.

Such lack of evolutionary innovation may be due, at least in part, to constraints placed by the limited surface area available for photosynthesis in the absence of thylakoids. With the emergence of thylakoids, new innovations on PSI structure and function were enabled. These include: 1) the gain of additional subunits, such as PsaK, PsaX and the PSI-associated IsiA-like Chl-binding proteins; 2) the emergence of subunit paralogs via gene duplication, facilitating the origin of adaptations such as photoacclimation to far-red light; and 3) the evolution of new oligomeric forms, such as PSI monomers and tetramers.

## Materials and Methods

### Strains and growth conditions

*A. panamensis* was isolated in a previous study (UTEX accession: 3164) (*13*). *G. violaceus* (also known as SAG 7.82 *G. violaceus*) was acquired from the Culture Collection of Algae at Göttingen University ("Sammlung von Algenkulturen der Universität Göttingen", SAG, Göttingen, Germany). *Synechocystis* 6803 was gifted by Dr. Hsiu-An Chu from Academia Sinica, Taiwan. The B-HEPES growth medium, a modified BG11 medium containing 1.1 g L^−1^ 4-(2-hydroxyethyl)-1-piperazine- ethanesulfonic acid (HEPES) pH 8.0 (adjusted with KOH), was used to cultivate the cultures, as previously described (*13, 23*). In a 30 °C growth chamber supplemented with 1 % (v/v) CO2 in the air, *A. panamensis* and *Synechocystis* 6803 cells were grown with cool white LED light at 10 and 50 μmol photons m^−2^ s^−1^, respectively (*13, 23*). Cool white LED light (5 μmol photons m^−2^ s^−1^) was used to grow *G. violaceus* in the air at 25 °C.

### Purification of trimeric Photosystem I complexes

PSI trimers from thylakoid-free strains (*A. panamensis* and *G. violaceus*) were purified as previously described with some modification (*11, 20, 51*). Cell pellets were resuspended in MES buffer (50 mM MES, pH 6.5, 10 mM CaCl2, and 10 mM MgCl2) and disrupted by glass beads (0.1 mm) in 2 mL screw cap tubes using a bead beater. The crude extract was centrifuged (2000 × *g* at 4 °C for 10 min) to remove the beads and unbroken cells.

Plasma membranes were recovered by centrifugation (13,800 × *g* at 4 °C for 10 min) after removal of beads and unbroken cells. In the dark, the plasma membrane fraction was incubated for 30 min at 4 °C in MES buffer containing 1 % (w/v) *n*-dodecyl-β-D-maltoside (β-DM). After centrifugation (13,800 × *g* at 4 °C for 10 min), the supernatant was layered on a linear sucrose density gradient (5–30 % (w/v) sucrose made with MES buffer containing 0.02 % (w/v) β-DM). The samples were centrifuged at 139,000 × *g* at 4 °C for 18 h. The lowest green band corresponding to PSI was collected and stored at −80 °C for further analysis.

PSI trimers from the thylakoid-containing strain *Synechocystis* 6803 were purified as described above with an additional step to collect thylakoid membranes. Cell pellets were resuspended in MES buffer (50 mM MES, pH 6.5, 10 mM CaCl2, and 10 mM MgCl2) and disrupted by a bead beater. After centrifugation (13,800 × *g* at 4 °C for 10 min) to remove beads and unbroken cells, the supernatant was further centrifuged (126,100 × *g* at 4 °C for 30 min) to collect thylakoid membranes. The membrane solubilization and ultracentrifugation steps were similar to the procedure for thylakoid-free cyanobacteria mentioned above.

### Absorption and low-temperature fluorescence spectroscopy

Absorption spectra of isolated PSI were measured using a Cary 60 UV-Vis spectrophotometer (Agilent, Santa Clara, CA, USA). Fluorescence emission spectra were obtained using a Hitachi F-7000 spectrofluorometer (Hitachi, Tokyo, Japan) with the excitation wavelength at 440 nm for Chl *a*. Liquid nitrogen was used to freeze the isolated cells or fractions to obtain 77 K fluorescence spectra.

### Pigment extraction and analysis

The pigment composition of PSI from *A. panamensis*, *G. violaceus*, and *Synechocystis* 6803 was determined by reversed-phase HPLC (*51*). Pigments were extracted from isolated PSI using acetone/methanol (1:1, v/v) as described previously (*42*). Extracts were centrifuged to remove insoluble proteins and cell debris, and the supernatant was collected and filtered through a 0.22-µm polytetrafluoroethylene membrane. A 100 µL aliquot was subjected to analysis by reversed-phase HPLC on a JASCO PU-4180 system equipped with a Discovery C18 column (4.6 mm × 25 cm). The gradient elution program used 100% methanol (solvent A) and 100% isopropanol (solvent B) with the following elution gradient [B, min]: [0%, 0 min], [97%, 75 min], [97%, 76 min], and [0%, 77 min] at a flow rate of 0.5 mL min^−1^. The elution was monitored at 270, 491, and 466 nm for quinones, carotenoids, and canthaxanthin, respectively. β-carotene, echinenone, and canthaxanthin ratios were calculated based on peak areas and extinction coefficients (141,000, 120,000, and 124,000 M^-1^ cm^-1^, respectively) (*52*).

### Polyacrylamide gel electrophoresis

Polyacrylamide gel electrophoresis in the presence of sodium dodecyl sulfate (SDS-PAGE) and urea (Tricine-urea-SDS-PAGE) was performed as previously described (*51, 53*). The samples were loaded onto a 16% (w/v) acrylamide containing 6 M urea gel.

After the separation of proteins by electrophoresis, the gel was stained with Coomassie brilliant blue G-250 to visualize all proteins. Selected subunits were identified by molecular weight and LC-MS/MS.

### In-solution and in-gel digestions for LC-MS/MS analysis

In-solution and in-gel digestions for LC-MS/MS analysis were performed as previously described (*23*). The protein solutions were diluted in 50 mM ammonium bicarbonate for in-solution digestions. They were subsequently reduced with 5 mM dithiothreitol at 60 °C for 45 min, followed by cysteine blocking with 10 mM iodoacetamide at 25 °C for 30 min. The samples were diluted with 25 mM ammonium bicarbonate and digested with sequencing-grade modified trypsin at 37 °C for 16 hours. The digested peptides were subjected to LC-MS/MS analysis.

The gel bands were dehydrated with 100% (v/v) acetonitrile by incubation at 37 °C for 30 min for in-gel digestions. The supernatant for each sample was discarded, and the samples were covered with 10 mM dithiothreitol in 100 mM ammonium bicarbonate at room temperature for 30 min. The supernatants were discarded, and the samples were treated with 50 mM iodoacetamide in 100 mM ammonium bicarbonate in the dark at room temperature for 30 min. The supernatant solutions were discarded, and the samples were washed three times with 100 mM ammonium bicarbonate. The samples were dehydrated at room temperature for 15 min using 100% (v/v) acetonitrile. The samples were air-dried at room temperature for 15 min, and the supernatants were discarded.

Trypsin digestion was conducted by incubating the samples overnight at 37 °C and covering them with sequencing-grade modified trypsin (0.01 μg μL^−1^) in 50 mM ammonium bicarbonate. The digested peptides were extracted by incubating them in acetonitrile containing 1 % (v/v) formic acid at 37 °C for 15 min. This extraction step was repeated twice, and the combined extracts were vacuum dried and analyzed by LC– MS/MS.

### Transmission electron microscopy of negatively stained PSI

*A. panamensis* PSI was buffer exchanged into MES buffer pH=6.5 with 0.02% β-DM. 4 µL of this sample at 2 µg Chl mL^−1^ was applied to a glow discharged (60 s, 25 mA) 400 mesh Cu grid with 5-6 nm formvar with 3-4 nm carbon (Electron Microscopy Sciences). The sample was negatively stained with 2% (w/v) uranyl acetate and imaged with a 120 kV Talos L120C TEM. Five micrograph images were collected. An example micrograph is shown in **fig. S6A**. The contrast transfer functions for the five micrographs were estimated with Ctffind-4.1.13 (*54*) within Relion 3.1 (*55*). 1,537 PSI particles were selected manually and 2D classification was performed (**fig. S6B**).

### Grid preparation for cryo-EM

A glow discharged (30 s, 25 mA) Quantifoil 2/1 Au 300 mesh electron microscopy grid (Electron Microscopy Sciences) was mounted in a Thermo Fisher Vitrobot Mark IV system set to 100% humidity and 4 °C. 3 µL of ∼2 mg Chl mL^−1^ *A. panamensis* PSI in MES buffer pH=6.5 with 0.02% β-DM was applied to the grid. It was blotted by the Vitrobot set to 4 °C for 3 s with blot force=0 and plunged into liquid ethane. The sample was transferred to liquid nitrogen for storage.

### Cryo-EM screening and data collection

Initial screening for cryo-EM was performed on a 200 kV Thermo Fisher Glacios TEM. Six micrograph movies were collected and processed in Relion 3.1 (*55*). An example micrograph is shown in **fig. S7A**. Motion correction, alignment, and dose-weighting was performed with MotionCor2 (*56*). The contrast transfer functions for the five micrographs were estimated with Ctffind-4.1.13 (*54*). 439 PSI particles were selected manually, and these were used for 2D classification (**fig. S7B**).

The same grid was subsequently imaged for high-resolution data collection using a 300 kV Thermo Fisher Titan Krios G2 TEM with a slit size of 15 eV at 105,000 × nominal magnification. The defocus range was −0.8 to −2.2 µm. The pixel size was 0.825 Å . The total dose was 50 e^−^ (Å)^−2^. EPU (Thermo Fisher) was used to collect 12,562 micrograph movies. An example micrograph is show in **fig. S8A**.

### High-resolution data processing

All data processing steps were performed in Relion 3.1 (*55*) Motion correction, alignment, and dose-weighting were performed with MotionCor2 (*56*) and the contrast transfer functions were estimated with Ctffind-4.1.13 (*54*). 529 particles were manually selected to create 2D templates for autopicking. 991,198 positions were selected by Autopicking. These were subjected to two rounds of 2D classification (example classes are shown in **fig. S8B**), yielding 669,002 particles. One round of 3D classification (an example class is shown in **fig. S8C**) yielded 403,080 particles that were the particles used in the final reconstruction. Rounds of contrast transfer function refinement and Bayesian Polishing were performed that provided a final map at 2.4 Å global resolution based on the Gold-standard Fourier Shell Correlation (0.143) cutoff criterion (*55, 57*). A scheme of the data processing workflow described here is shown in **fig. S8D**.

### Model building

An initial model was created by generating homology models of each subunit with SwissModel (*58*). These were superimposed onto the corresponding subunits of the *T. vestitus* PSI structure (PDB 1JB0) (*4*) and combined into a single coordinate file with the cofactors extracted from 1JB0. This model was fit into the cryo-EM map using UCSF Chimera (*59*). Coot (*60*) was used for manually editing the structure. Automated refinement was performed using real_space_refine (*61*) in Phenix (*62*).

### Phylogenetic analysis

Amino acid sequences for PSI subunits were downloaded from the NCBI database on the 17th of May 2024 (PsaA, PsaB, PsaC, PsaD, PsaE, PsaL, and PsaM) and on the 27th of August 2024 (PsaF, PsaI, and PsaJ) using PSI-BLAST. Sequence redundancy was reduced to 98% sequence identity for all subunits except PsaC, PsaL and PsaK. For PsaC no redundance filter was applied. For PsaL and PsaK a 95% cut off was applied.

Singleton sequences with large insertions or deletions were removed. The C-terminal extension of PsaB in the sequences from *G. violaceus* and *G. morelensis* was also trimmed. Alignments were carried out with Clustal Omega (*63*), using 5 combined guided trees and HMM iterations. Maximum Likelihood tree inference was performed for each individual subunit with IQ-TREE 2.2 (*64*). The best fitting substitution model was calculated automatically by the software and branch support values were calculated with both ultrafast bootstrap until the correlation coefficient converged, and additionally, with the average likelihood ratio test. An additional phylogenetic tree was inferred using a combined sequence alignment of PsaA and PsaB, activating the ancestral sequence reconstruction feature (-asr) in IQ-TREE. Furthermore, an additional sequence alignment of combined PsaA and PsaB sequences was prepared by removing poorly aligned regions using the tool Gblocks, as implemented in the program Seaview 5.0.5 and enabling options for a less stringent selection of conserved sites (*65*). All sequence datasets and phylogenies are presented in the Supplementary Data.

## Notes

### Competing Interest Statement

The authors have declared no competing interest.

