## Supplementary Materials for "Structure and evolution of Photosystem I in the early-branching cyanobacterium *Anthocerotibacter panamensis*"

**Affiliations**

### This PDF file includes:

Figs. S1 to S17

Tables S1 to S5

Data S1

### Supplementary Figures

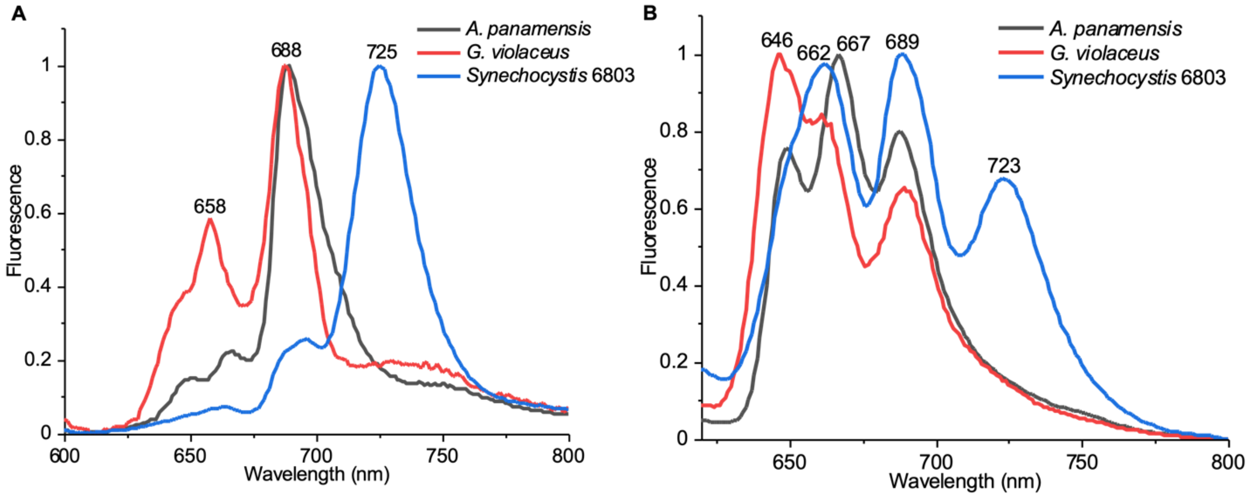

**Fig. S1. Low temperature (77 K) fluorescence emission spectrum of *A. panamensis, G. violaceus*, and *Synechocystis* 6803 cells**. (**A**) Excitation wavelength at 440 nm. (**B**) Excitation wavelength at 580 nm.

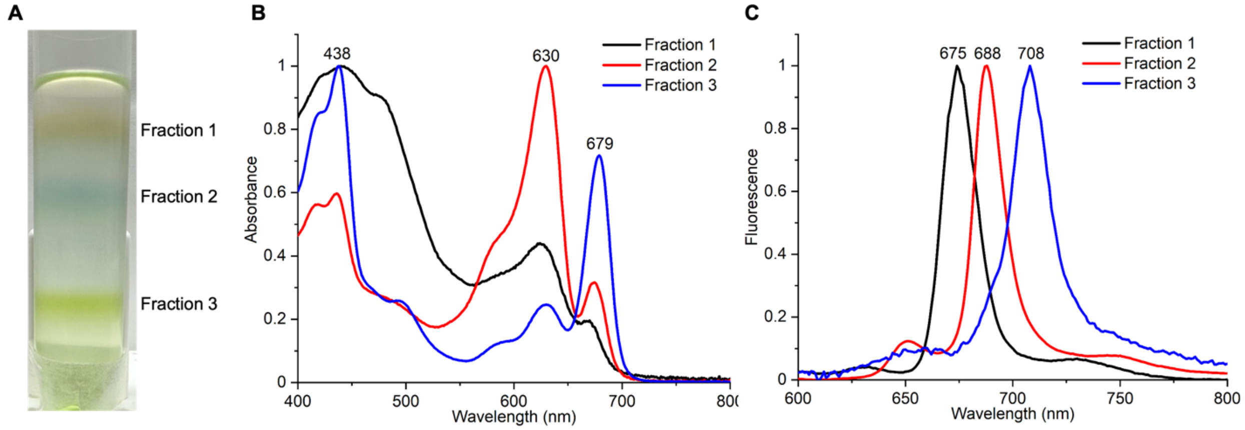

**Fig. S2. Spectroscopic characterization of the PSI fractions from *A. panamensis*.** (**A**) Separation of solubilized *A. panamensis* cell membranes by sucrose density gradient centrifugation*.* (**B**) Room-temperature absorption spectrum of sucrose gradient fractions. (**C**) 77 K fluorescence emission spectrum of sucrose gradient fractions excited at 440 nm.

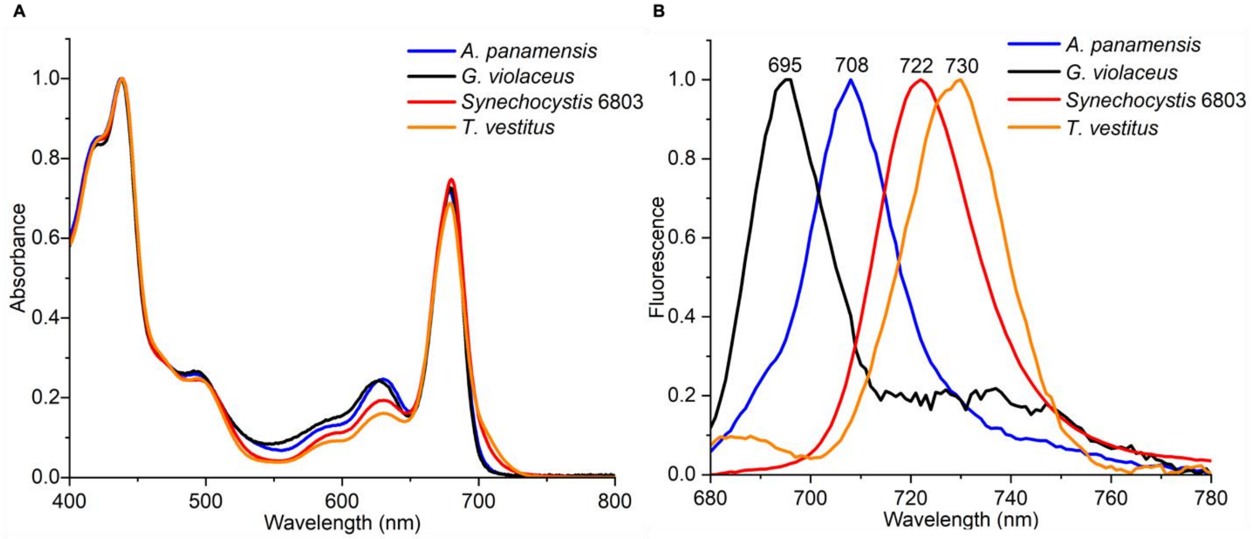

**Fig. S3. Absorption spectra and 77 K fluorescence emission of PSI from *A. panamensis* (blue), *G. violaceus* (black), *Synechocystis* 6803 (red), and *T. vestitus* (orange).** (**A**) Absorption spectra measured at room-temperature and normalized by their maximum peak intensities. (**B**) Fluorescence spectra measured at 77 K and normalized by their maximum peak intensities. The absorption spectra data and 77 K fluorescence-emission data of *T. vestitus* were adapted from Çoruh et al., 2021 (*25*).

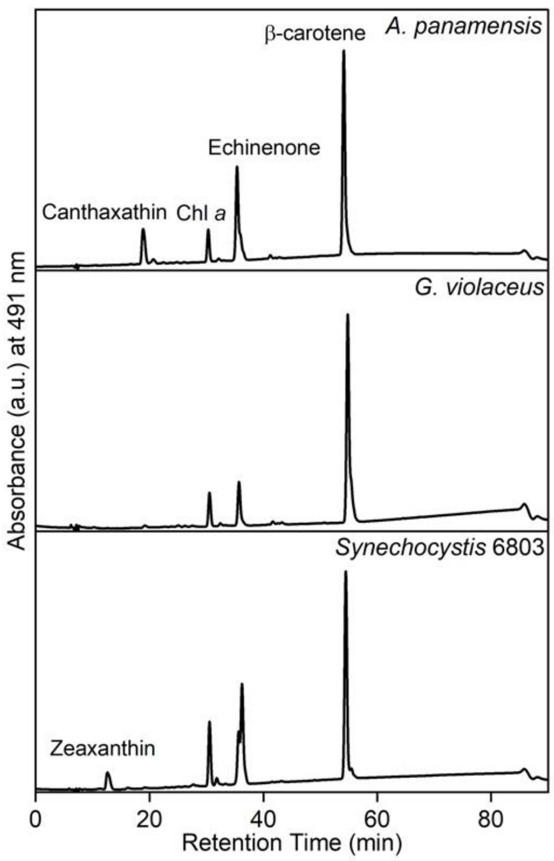

**Fig. S4. HPLC analysis of PSI from *A. panamensis*, *G. violaceus*, and *Synechocystis* 6803.** Four major pigment peaks are eluted from the PSI of *A. panamensis* and are identified as canthaxanthin, chlorophyll *a* (Chl *a*), echinenone, and β-carotene, respectively, based on their elution time and characteristic absorption spectra (**Fig. S5**).

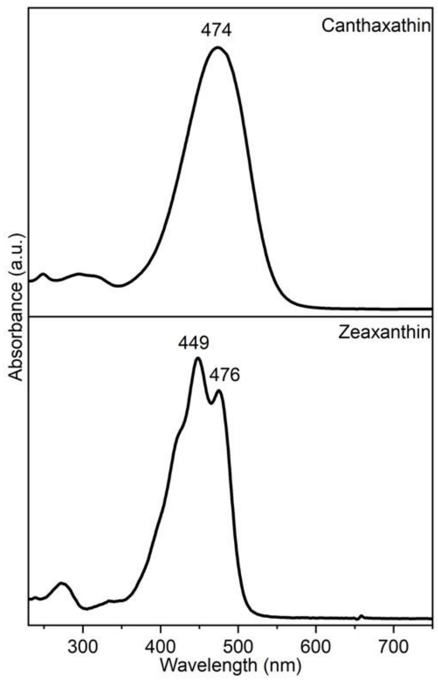

**Fig. S5. The absorption spectra of components eluted in HPLC analysis. (A)** Canthaxanthin in *A. panamensis*. **(B)** Zeaxanthin in *Synechocystis* 6803.

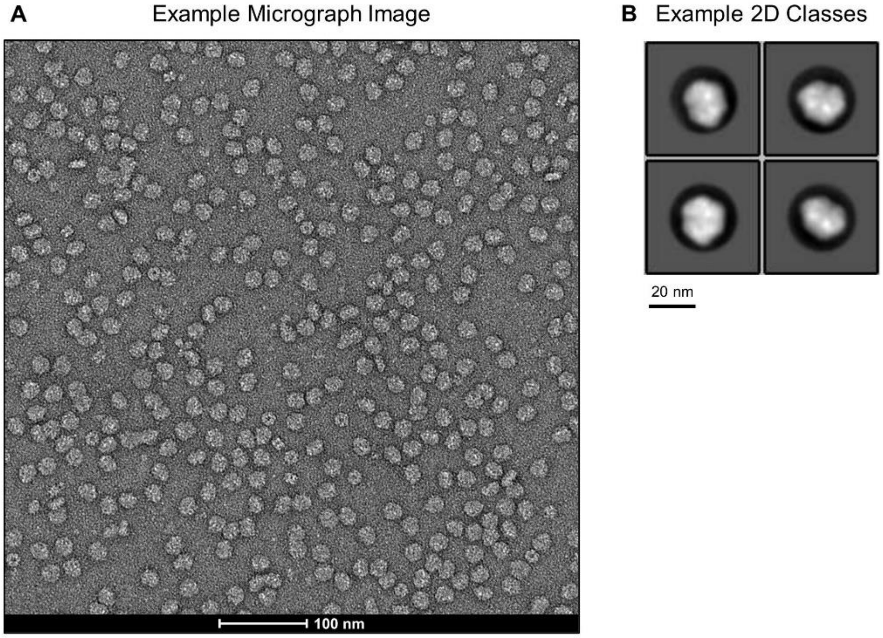

**Fig. S6. TEM of negatively stained *A. panamensis* PSI.** (**A**) Example micrograph. (**B**) Example 2D Classes.

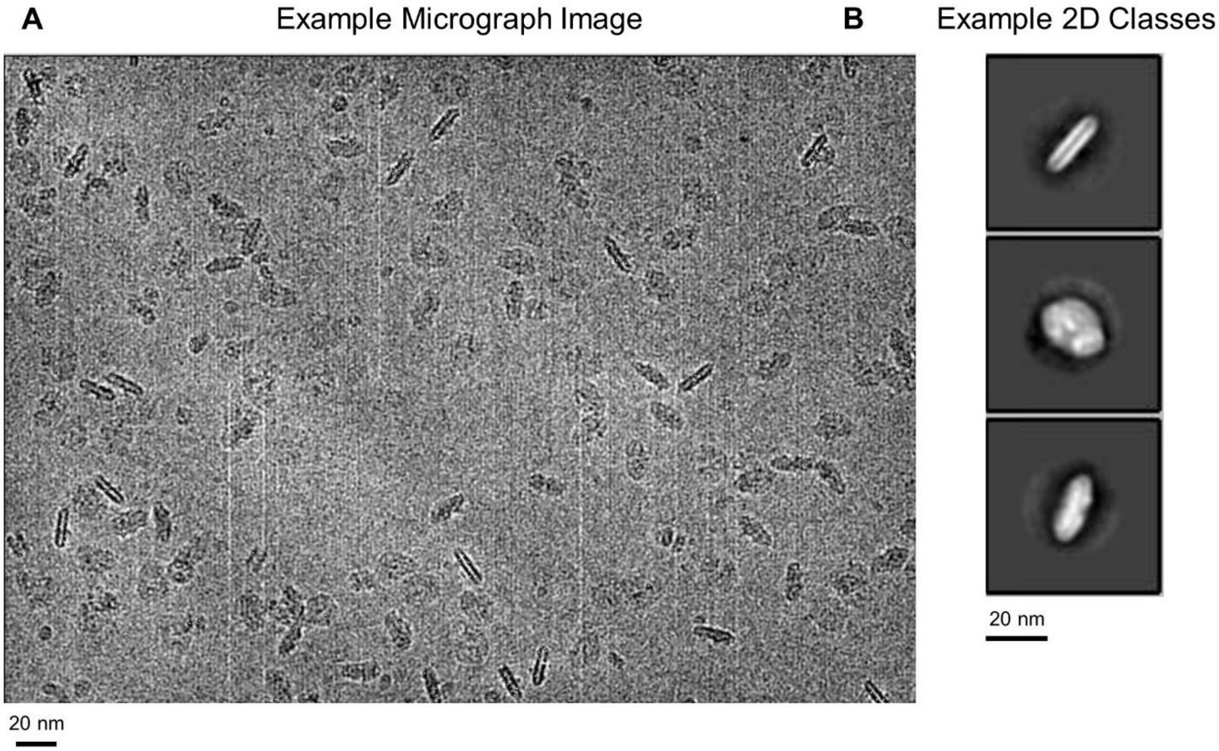

**Fig. S7. Glacios cryo-EM screening of *A. panamensis* PSI.** (**A**) Example micrograph. (**B**) Example 2D Classes.

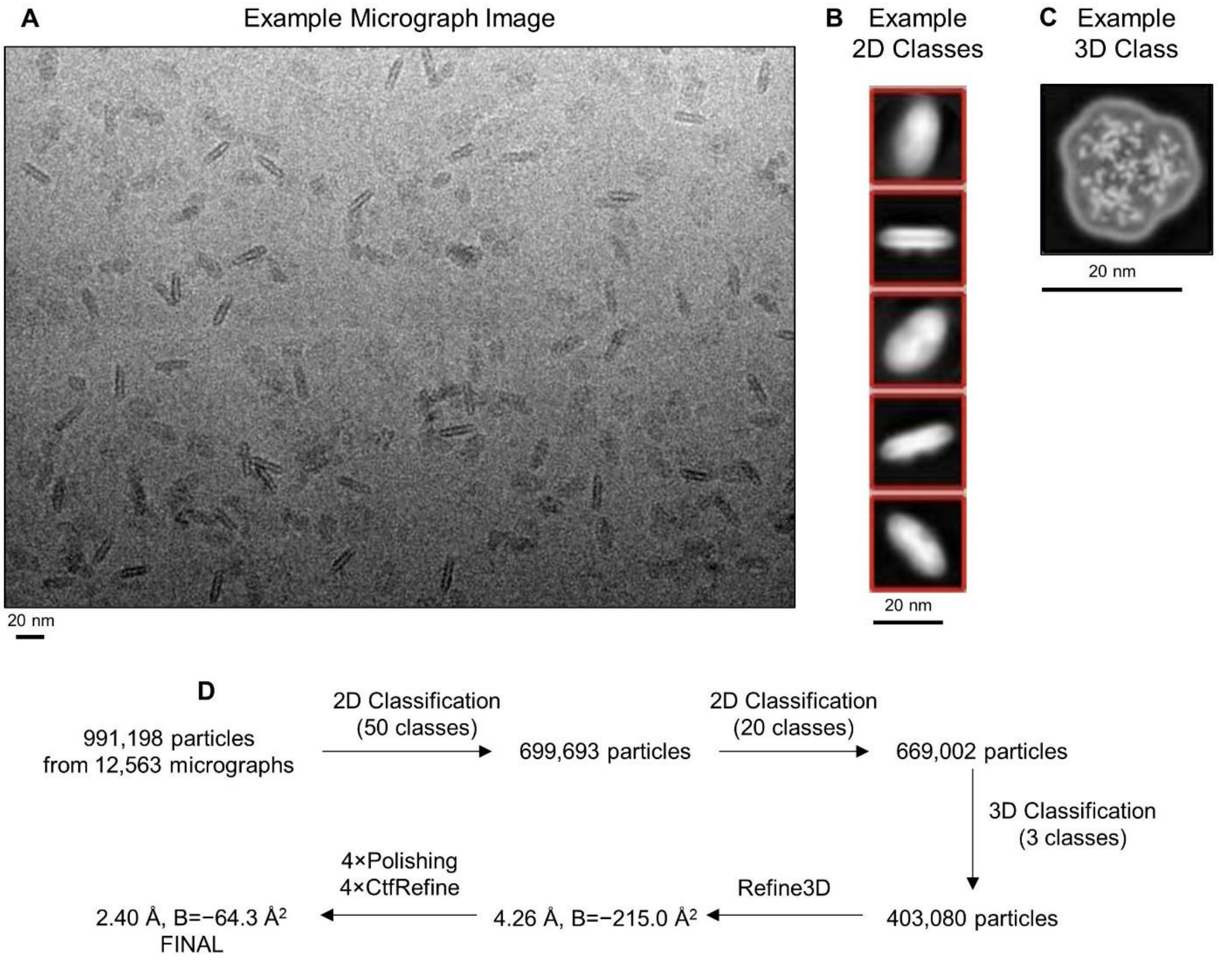

**Fig. S8. Krios cryo-EM screening of *A. panamensis* PSI.** (**A**) Example micrograph. (**B**) Example 2D Classes. (**C**) Example slice through a 3D class. (**D**) Overall workflow of data processing.

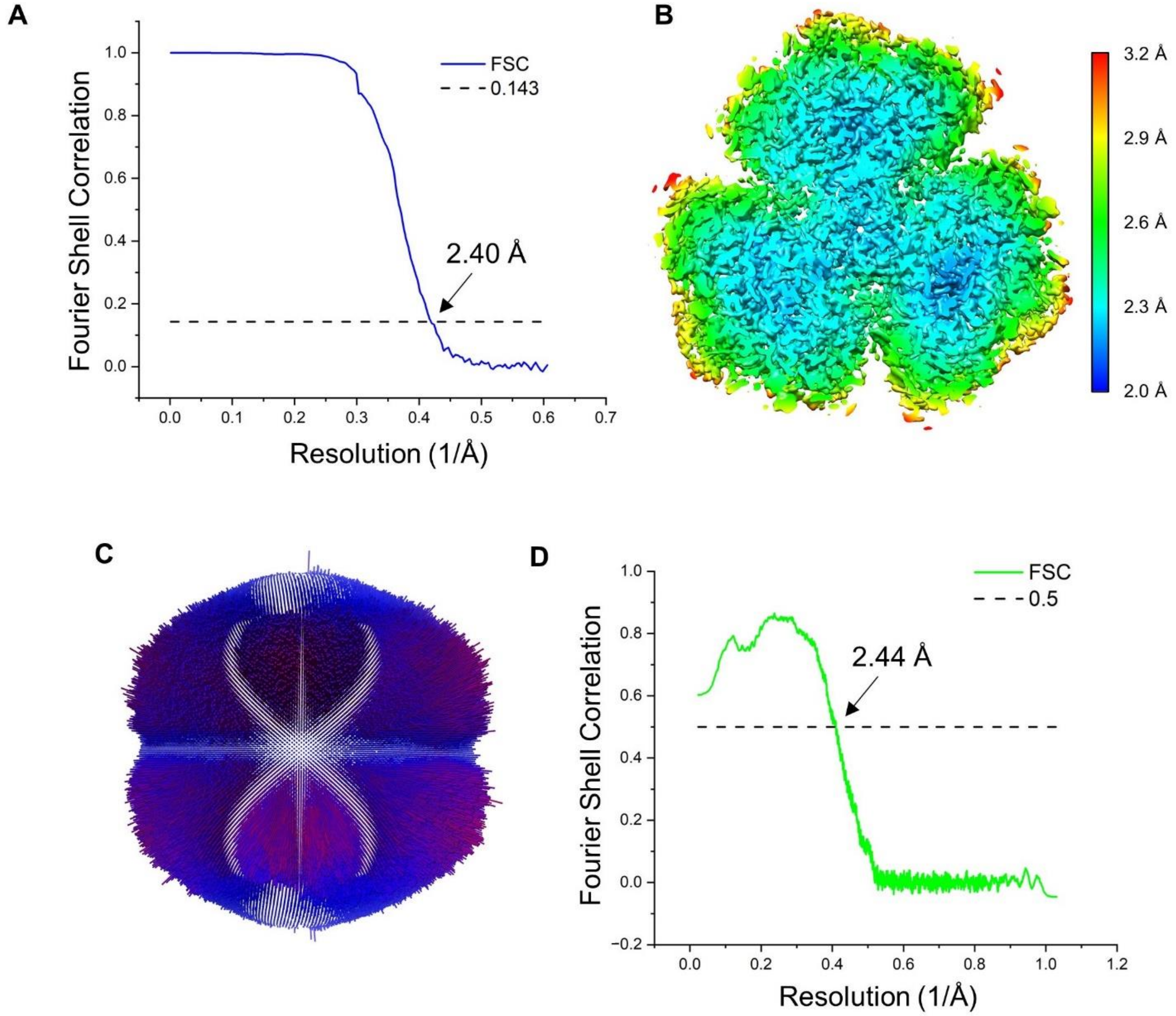

**Fig. S9. Resolution and completeness of cryo-EM data.** (**A**) Map-to-map Fourier Shell Correlation. (**B**) Local resolution map. (**C**) Angular distribution of the cryo-EM data. (**D**) Map-to-model Fourier Shell Correlation.

Phylloquinone-4

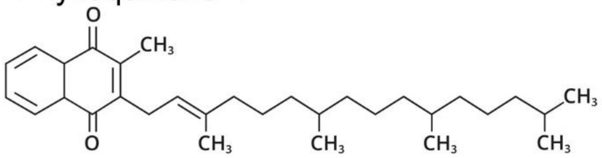

Menaquinone-4

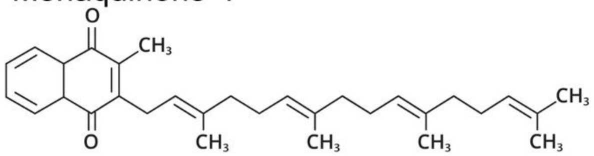

**Fig. S10. Structures of PhQ-4 and MQ-4.**

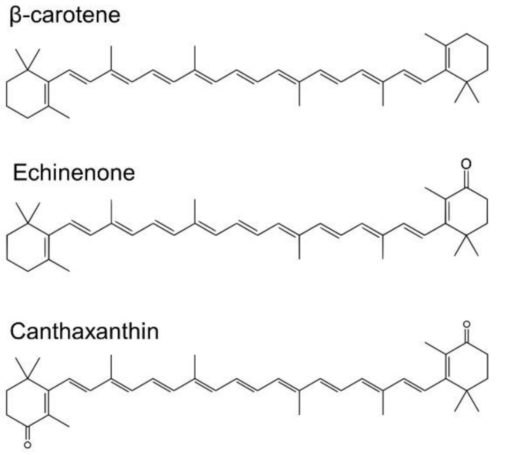

**Fig. S11. Structures of β-carotene, echinenone, and canthaxanthin.**

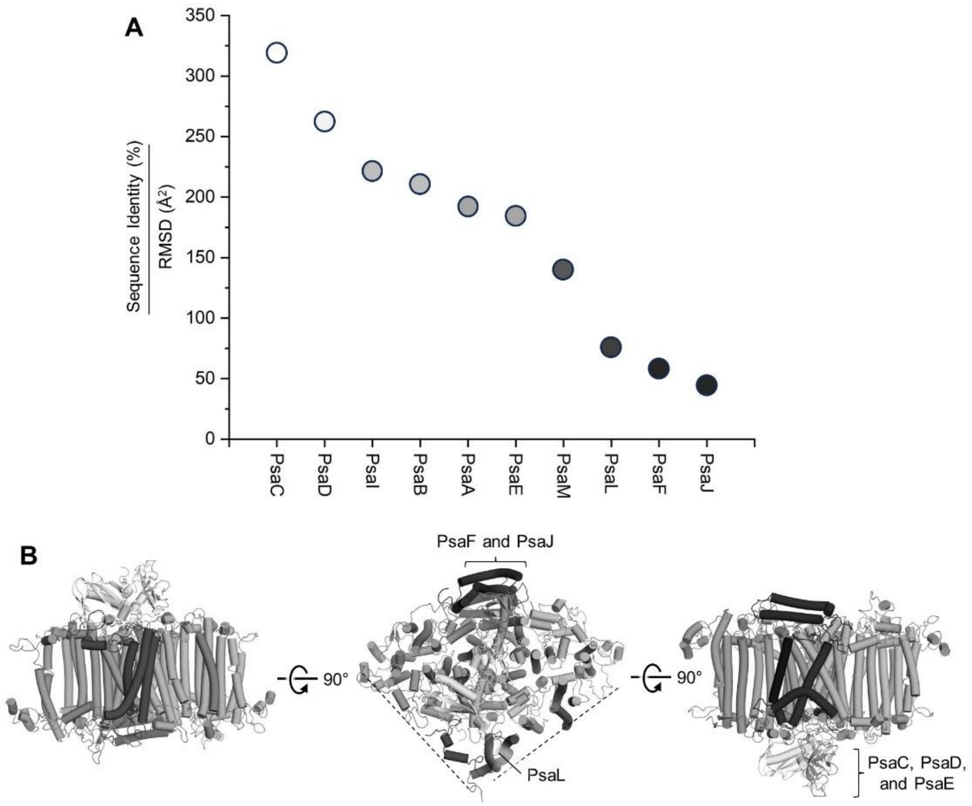

**Fig. S12. Comparison of the *A. panamensis* PSI structure to *G. violaceus* PSI.** (**A**) Plot of sequence identity (%) divided by RMSD ($Å^2$) for each subunit. (**B**) Structure of an *A. panamensis* PSI monomer where subunits are colored as the shades of circles in panel a. In the center panel, dashed lines correspond to monomer-monomer interfaces. Note that this table is generated from the data shown in **Table S3**.

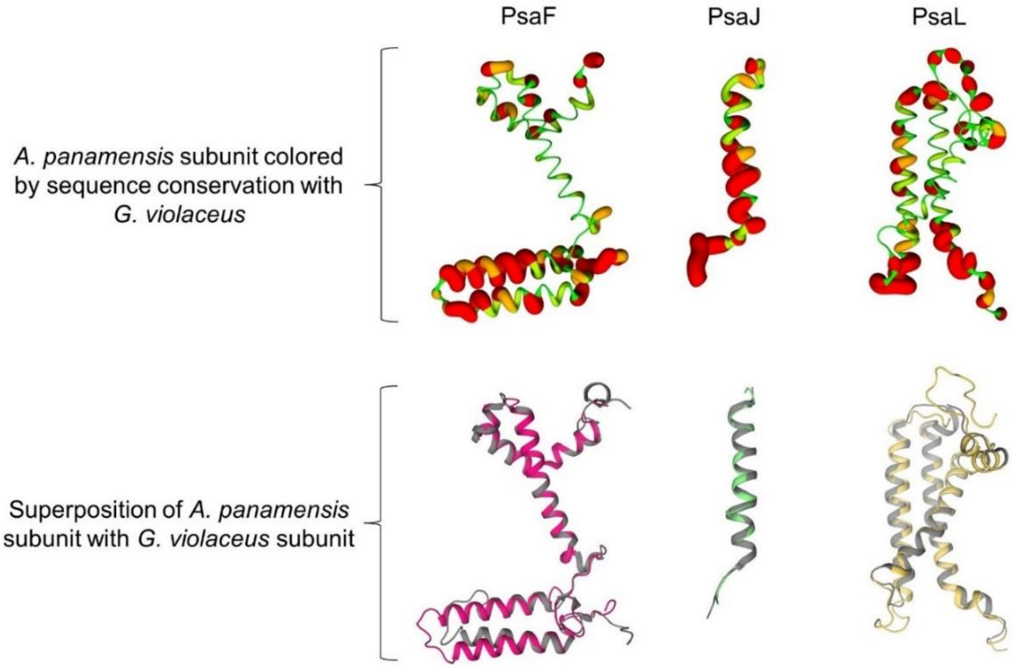

**Fig. S13. Structural view of sequence conservation and superposition of selected *A. panamensis* PSI subunits with *G. violaceus* subunits.**

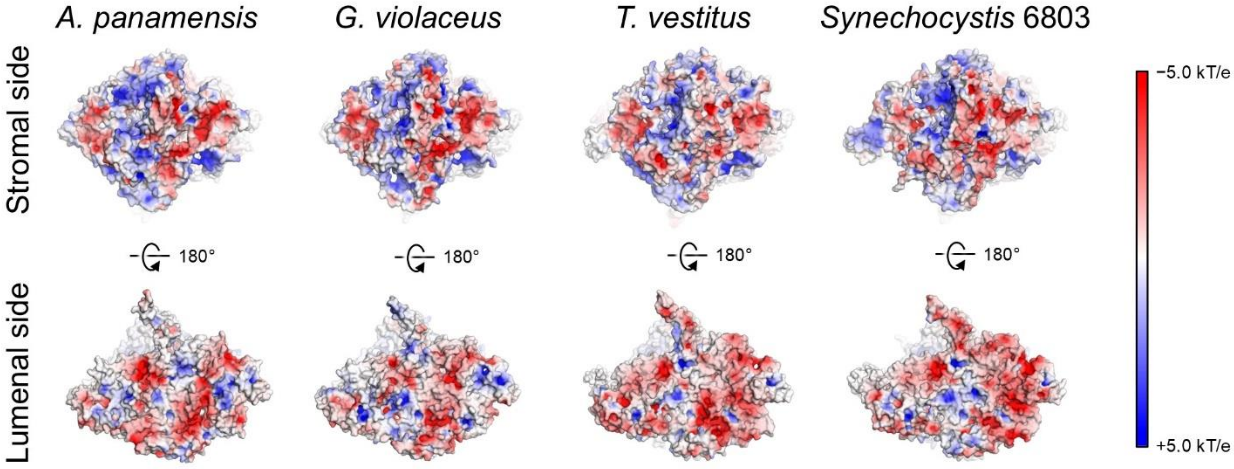

**Fig. S14. Surface electrostatics comparison between selected PSI complexes.** Note that the lumenal side of the PsaL region is more negatively charged in *T. vestitus* and *Synechocystis* 6803 compared to *A. panamensis* and *G. violaceus*.

**A**

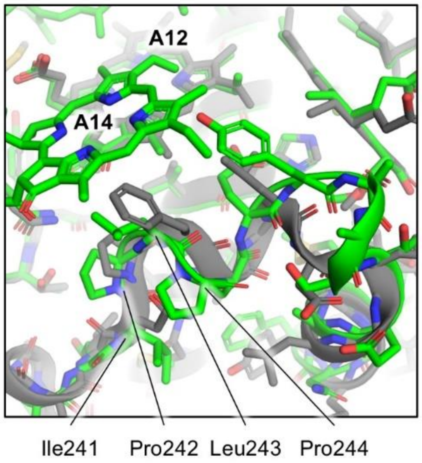

**B**

| | | |
|---|---|---|
| Ancestral PsaA | DIPLPHEFI | |
| PsaA *T. vestitus* | DIPLPHEFI 243 | (has A14) |
| PsaA *Synechocystis* 6803 | DIPLPHEFI 243 | (has A14) |
| PsaA *A. panamensis* | DIPLPHEYA 248 | (has A14) |
| PsaA *C. vandensis* | AIPLPHEYA 247 | |
| PsaA ES-bin-313 | EVVIP---G 244 | |
| PsaA ES-bin-141 | DVNPFL--G 256 | |
| PsaA *G. violaceus* | QVNPFA--G 245 | (lacks A14) |
| PsaA *G. morelensis* | QVNAFA--G 245 | |
| PsaA *G. kilaueensis* | QVNPFL--G 345 | |

**Fig. S15. Structural and sequence analysis of the A14/A12 Chl site.** (**A**) Superposition of the *A. panamensis* and *G. violaceus* PSI structures in the vicinity of the A14/A12 Chl site. Residues conserved in *A. panamensis*, *T. vestitus*, and *Synechocystis* 6803 PSI are labeled. (**B**) Partial sequence alignment of PsaA in the vicinity of A14/A12. The Clustal Omega sequence identifiers are shown at the bottom of each position (*59*).

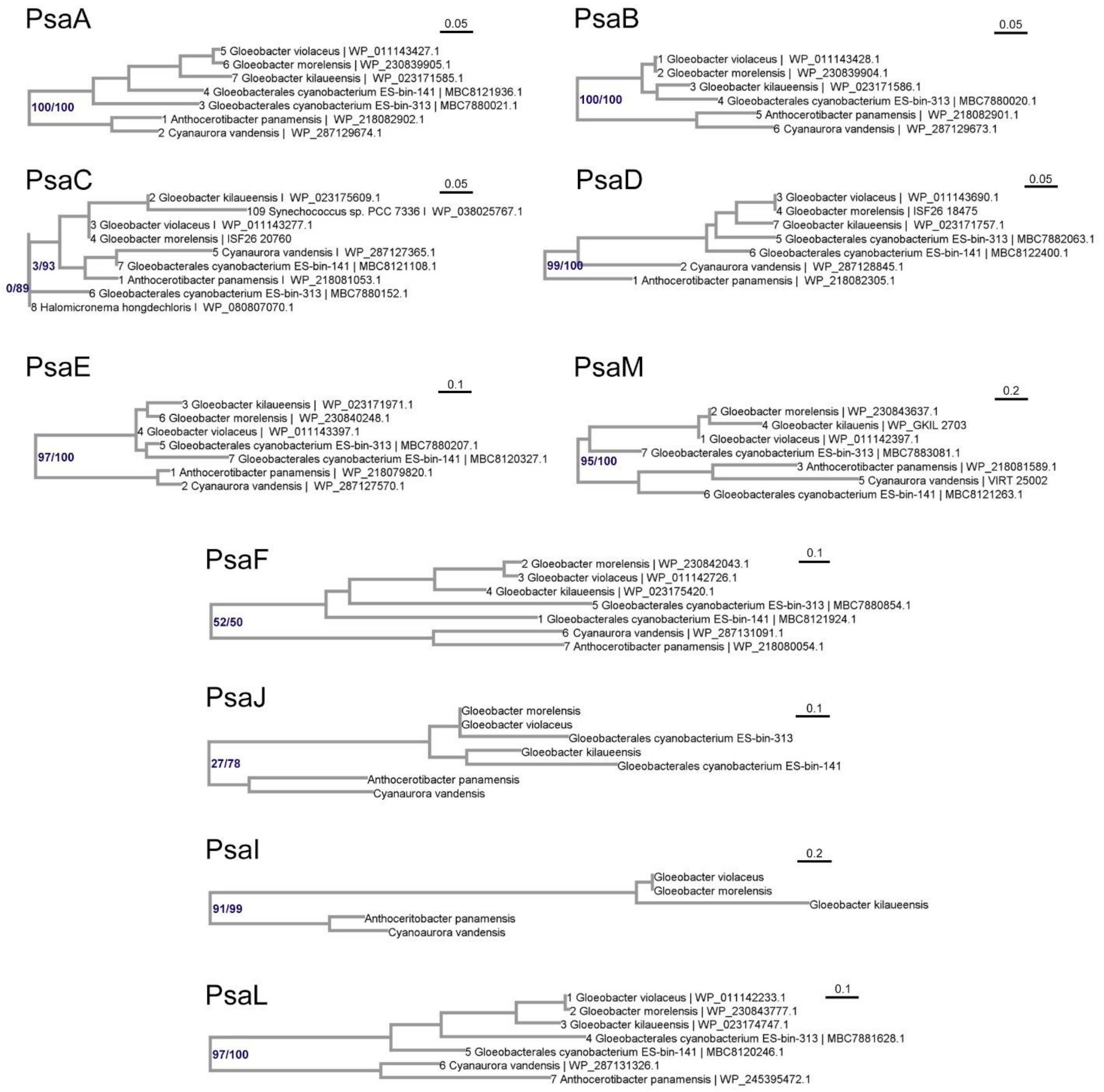

**Fig. S16. Detail of PSI subunit evolution for *A. panamensis* and close relatives.** Maximum Likelihood phylogenetic trees of each PSI subunit in *A. panamensis.* Trees were run as described in **Materials and Methods** and rooted at the branching point of the Gloeobacterales. Only Gloeobacterales are shown here. The trees show that PSI subunits in Gloeobacterales show strong congruence across subunits, suggesting that these have been passed down vertically since their most recent common ancestor. No evidence for replacement of subunits via horizontal gene transfer or gain of subunit paralogs via duplication is observed. Full trees and sequence alignments are provided as **Data S1**. Scale bars represent amino acids substitutions per site.

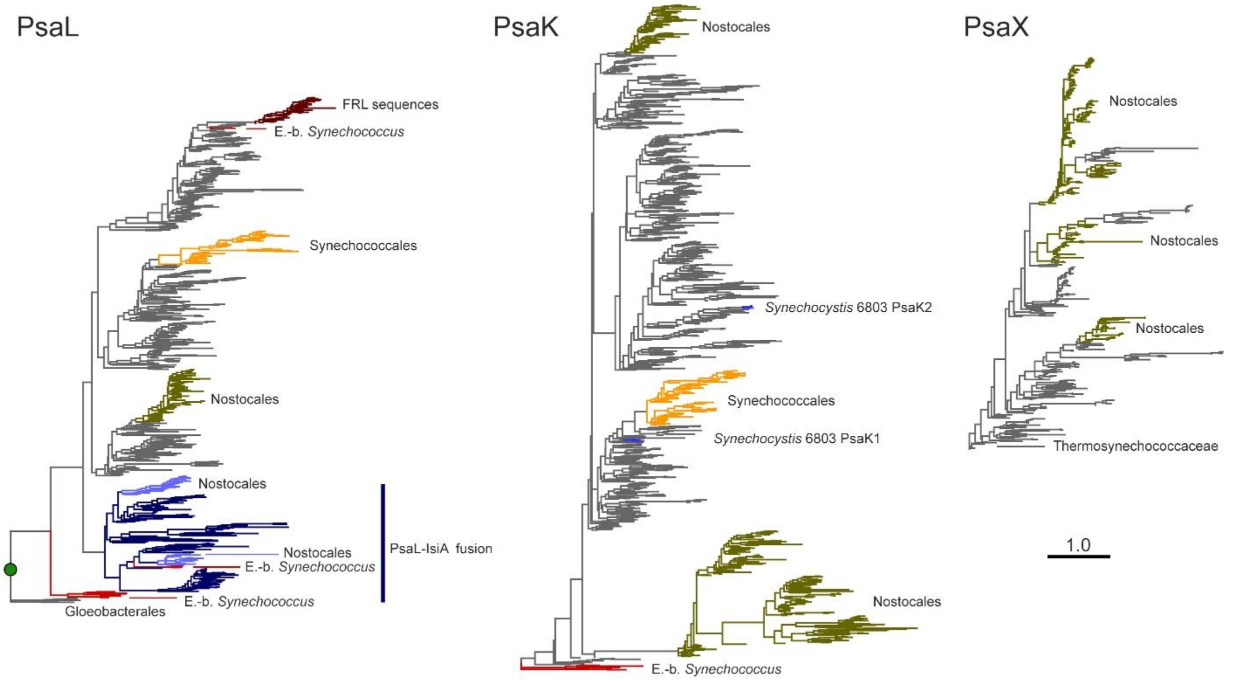

**Fig. S17. Maximum Likelihood tree of PsaL, PsaK, and PsaX.** The PsaL phylogeny was rooted at the branching point of the Gloeobacterales. Several gene duplications are noted, some of which appear to be very ancient. In particular, that leading to the PsaL-IsiA fusion, which likely occurred after the early divergence of the clade containing the early-branching *Synechococcus* (E.-b *Synechococcus*). The PsaL-IsiA fusion appears to have duplicated at least once as evidenced by the presence of two distinct paralogues in Nostocales (e.g. heterocystous cyanobacteria). Synechococcales denotes the *Synechococcus*-*Prochlorococcus* clade. FRL sequences represent another duplication leading to the paralogue found in the far-red light photoacclimation gene cluster. PsaK is absent in Gloeobacterales but it is found in most other cyanobacteria, including the early-branching *Synechococcus*, where the tree was rooted for display. Several duplications are noted as evidenced by the two paralogues found in Nostocales as well as the two paralogues known in the model cyanobacterium *Synechocystis* 6803. In contrast, PsaX has a limited distribution. It is neither found in any of the most basal cyanobacterial groups nor the Synechococcales, except for a few strains in the *Thermosynechococcus* group, where the tree was rooted for display. The subunit appears to be most widely distributed in Nostocales and their relatives, as well as other macrocyanobacteria.

### Supplementary Tables

**Table S1. Mass spectrometric analysis of the *A. panamensis* PSI fraction from sucrose gradient.** The criteria for identifying a protein were two peptides and a false discovery rate below 0.01. The protein scores are calculated as the sum of the scores of peptides.

| Protein | Score | Coverage (%) | Molecular weight (kDa) |
|---|---|---|---|
| PsaA | 5,463 | 59 | 86.6 |
| PsaB | 4,590 | 51.8 | 82.8 |
| PsaC | 1,271 | 98.8 | 8.8 |
| PsaD | 3,908 | 95.8 | 15.5 |
| PsaE | 2,418 | 98.4 | 7.2 |
| PsaF | 2,669 | 74 | 19.5 |
| PsaL | 1,175 | 96.3 | 17 |
| Heat shock protein 60 kDa family chaperone GroEL | 7,032 | 87.9 | 57.8 |
| Phycobilisome rod linker polypeptide, phycocyanin-associated (CpcN) | 5,107 | 60.7 | 133.8 |
| Phycocyanin beta chain | 5,060 | 100 | 18.4 |
| Phycocyanin alpha chain | 4,742 | 99.4 | 17.8 |
| Heat shock protein 60 kDa family chaperone GroEL | 3,902 | 71.2 | 58.5 |
| Glutamine synthetase type I (EC 6.3.1.2) | 3,344 | 94.5 | 52.4 |
| ATP synthase beta chain (EC 3.6.3.14) | 2,331 | 78.5 | 51.9 |
| Ribulose bisphosphate carboxylase large chain (EC 4.1.1.39) | 1,616 | 61.8 | 52.7 |
| ATP synthase alpha chain (EC 3.6.3.14) | 1,482 | 64.1 | 54 |
| CRISPR-associated protein, Cse4 family | 1,397 | 58.2 | 41.4 |
| hypothetical protein | 1,354 | 46.1 | 88.9 |
| Photosystem II CP47 protein (PsbB) | 1,265 | 38.9 | 59.7 |
| Protein QmcA (possibly involved in integral membrane quality control) | 1,165 | 52.2 | 35.7 |
| Phycobilisome phycoerythrin-associated linker polypeptide CpcJ | 968 | 26.6 | 60.2 |
| Branched-chain amino acid ABC transporter, substrate-binding protein LivJ (TC 3.A.1.4.1) | 685 | 52.7 | 40.4 |
| Heat shock protein 10 kDa family chaperone GroES | 651 | 95.1 | 11 |

**Table S2. Mass spectrometric identification of in-gel digested proteins from SDS-PAGE.** Ten selected bands from the SDS-PAGE of PSI, labeled 1–7 in **Fig. 1e** were analyzed. The protein scores are calculated as the sum of the scores of peptides. The top four proteins with the highest scores are included. The high-molecular-weight Psa subunits detected at Band 6-7 are likely degraded proteins, indicated by their low scores and coverages. The identified PSI subunits are highlighted in green.

| | Protein | Score | Coverage (%) | Molecular weight (kDa) |
|---|---|---|---|---|
| **Band 1** | PsaB | 1544 | 25.9 | 82.8 |
| | PsaA | 1499 | 32.5 | 86.6 |
| | CpcN | 1446 | 32.5 | 133.8 |
| | hypothetical protein | 1308 | 61.7 | 61.1 |
| **Band 2** | CpcN | 484 | 25.5 | 133.8 |
| | CpcJ | 425 | 42.2 | 60.2 |
| | SSU ribosomal protein S4p | 401 | 63.6 | 23.5 |
| | PsaB | 228 | 14.6 | 82.8 |
| **Band 3** | PsaD | 1585 | 99.3 | 15.5 |
| | PsaF | 1478 | 45.8 | 19.5 |
| | Phycocyanin alpha chain | 742 | 84 | 17.8 |
| | Phycocyanin beta chain | 736 | 83.1 | 18.4 |
| **Band 4** | PsaD | 292 | 86.7 | 15.5 |
| | PsaC | 234 | 74.1 | 8.8 |
| | PsaL | 224 | 31.3 | 17 |
| | SSU ribosomal protein S17p | 174 | 45.1 | 11.5 |
| **Band 5** | PsaC | 666 | 100 | 8.8 |
| | PsaE | 349 | 98.4 | 7.2 |
| | PsaF | 123 | 21.5 | 19.5 |
| | Phycocyanin alpha chain | 86 | 30.1 | 17.8 |
| **Band 6** | Ribulose bisphosphate carboxylase large chain | 69 | 4 | 52.7 |
| | PsaA | 52 | 2.3 | 86.6 |
| | PsaC | 45 | 35.8 | 8.8 |
| | ABC transporter, RND-adapter-like protein | 36 | 1.4 | 46.7 |
| **Band 7** | PsaB | 81 | 1.7 | 82.8 |
| | Ribulose bisphosphate carboxylase large chain | 59 | 5.5 | 52.7 |
| | Alanyl-tRNA synthetase | 39 | 0.7 | 94.2 |
| | PsaA | 37 | 3.2 | 86.6 |

**Table S3. Cryo-EM data collection, refinement, and validation statistics for *A. panamensis* PSI.**

| | |
|---|---|
| **Data collection and processing** | |
| Magnification | x105,000 |
| Voltage (kV) | 300 |
| Electron exposure ($e^{-}Å^{-2}$) | 50.0 |
| Defocus range (μm) | −0.8 to −2.2 |
| Pixel size (Å) | 0.825 |
| Symmetry imposed | C3 |
| Initial particle images (no.) | 991,198 |
| Final particle images (no.) | 403,080 |
| Map resolution (Å) | 2.40 |
| FSC threshold | 0.143 |
| **Refinement** | |
| Initial model used (PDB code) | 7F4V |
| Model resolution (Å) | 2.44 |
| FSC threshold | 0.5 |
| Map resolution range (Å) | 2.10-3.20 |
| Map-sharpening *B* factor ($Å^2$) | −64.3 |
| Model composition | |
| Non-hydrogen atoms | 69,921 |
| Protein residues | 6,498 |
| Ligands | 366 |
| *B* factors ($Å^2$) | |
| Protein | 20 |
| Ligands | 23 |
| R.m.s. deviations | |
| Bond lengths (Å) | 0.009 |
| Bond angles (°) | 2.288 |
| **Validation** | |
| MolProbity | 2.48 |
| Clashscore | 10.63 |
| Rotamer outliers (%) | 4.62 |
| Ramachandran plot | |
| Favored (%) | 93.40 |
| Allowed (%) | 5.78 |
| Disallowed (%) | 0.82 |

**Table S4. Cryo-EM model composition (per monomer).**

| | **Number** |
|---|---|
| **Protein subunits** | 10 |
| **Chlorophyll *a*** | 88 |
| **Chlorophyll *a′*** | 1 |
| **Menaquinone-4** | 2 |
| **β-carotene** | 22 |
| **Echinenone** | 1 |
| **[4Fe-4S] cluster** | 3 |
| **Phosphatidylglycerol** | 3 |
| **Distearoyl-monogalactosyl-diglyceride** | 1 |
| ***n*-dodecyl β-D-maltoside** | 1 |

**Table S5. Sequence identities and root-mean square deviation (RMSD) of *A. panamensis* PSI subunits with those from other selected cyanobacterial species for which structures are available.** Values are colored from green to white corresponding to most to least similar.

| | Sequence Identity (%) | RMSD (Å) | | Sequence Identity (%) | RMSD (Å) |
|---|---|---|---|---|---|
| **PsaA** | | | **PsaF** | | |
| *G. violaceus* | 76.74 | 0.400 | *G. violaceus* | 36.90 | 0.634 |
| *T. vestitus* | 73.73 | 0.501 | *T. vestitus* | 39.07 | 0.881 |
| *Synechocystis* 6803 | 72.76 | 0.441 | *Synechocystis* 6803 | 35.53 | 0.981 |
| **PsaB** | | | **PsaI** | | |
| *G. violaceus* | 79.92 | 0.380 | *G. violaceus* (PsaZ) | 53.12 | 0.240 |
| *T. vestitus* | 74.07 | 0.454 | *T. vestitus* | 29.03 | 0.662 |
| *Synechocystis* 6803 | 76.56 | 0.438 | *Synechocystis* 6803 | 31.25 | 0.742 |
| **PsaC** | | | **PsaJ** | | |
| *G. violaceus* | 96.30 | 0.302 | *G. violaceus* | 17.50 | 0.395 |
| *T. vestitus* | 92.59 | 0.362 | *T. vestitus* | 28.21 | 1.086 |
| *Synechocystis* 6803 | 93.83 | 0.356 | *Synechocystis* 6803 | 26.32 | 1.064 |
| **PsaD** | | | **PsaL** | | |
| *G. violaceus* | 79.72 | 0.304 | *G. violaceus* | 54.42 | 0.717 |
| *T. vestitus* | 57.97 | 0.486 | *T. vestitus* | 42.18 | 2.246 |
| *Synechocystis* 6803 | 58.70 | 0.452 | *Synechocystis* 6803 | 43.92 | 0.993 |
| **PsaE** | | | **PsaM** | | |
| *G. violaceus* | 75.00 | 0.407 | *G. violaceus* | 58.06 | 0.415 |
| *T. vestitus* | 61.90 | 0.400 | *T. vestitus* | 38.71 | 0.798 |
| *Synechocystis* 6803 | 58.73 | 0.330 | *Synechocystis* 6803 | 38.71 | 0.842 |

**Data S1. Phylogenetic trees and sequence alignments used for Fig. S16.** (external)
